# Biodiversity material remains where capacity and governance are strong, but taxonomic resources concentrate with geopolitical power

**DOI:** 10.64898/2026.06.23.734087

**Authors:** Mario R. Moura, Ricardo Henrique Pereira da Silva, Mariana Pedrozo, Jhonny J. M. Guedes, Peter Uetz, Matheus de T. Moroti

**Affiliations:** Departamento de Sistemática e Ecologia, Universidade Federal da Paraíba, 58051-900, João Pessoa, Brazil; Programa de Pós-graduação em Biodiversidade, Universidade Federal da Paraíba, 58397-000, Areia, Brazil; Instituto de Biociências, Universidade Federal de Mato Grosso do Sul, 79070-900, Campo Grande, Brazil; Departamento de Biologia Animal, Universidade Estadual de Campinas, 13083-862, Campinas, Brazil; Department of Biogeography and Global Change, Museo Nacional de Ciencias Naturales (MNCN-CSIC), Madrid, Spain; Center for Biological Data Science, Virginia Commonwealth University, Richmond, VA, USA

**Keywords:** Biodiversity documentation, knowledge inequities, species discovery bias, scientific colonialism, type specimens

## Abstract

**Aim:** Biodiversity-rich regions often lack the scientific infrastructure needed to document and curate their own biodiversity, creating inequalities in access to taxonomic reference material. We investigated how biological, institutional, and geopolitical factors shape the retention, extraction, and appropriation of reptile holotypes, the name-bearing specimens upon which species descriptions are based.

**Location:** Global.

**Taxon:** Reptiles.

**Methods:** We compiled a historical dataset of reptile holotype origins and destinations spanning 1758–2024 to reconstruct long-term patterns of retention and international specimen flows. We then quantified species-level holotype retention, holotype flows between country pairs, and country-level patterns of retention, appropriation, and network centrality for the period 1990–2024, and used generalised linear mixed models to assess the biological, institutional, and geopolitical determinants of these contemporary circulation processes.

**Results:** Although nearly 90% of reptile species described originated in the Global South, less than a quarter of their holotypes remain housed there. Historically, exported holotypes consistently outnumbered retained holotypes on a decadal basis until the early twenty-first century. Retention was promoted by local scientific capacity, institutional infrastructure, collector involvement in species descriptions, and environmental governance. In contrast, extraction was concentrated in highly endemic regions with limited scientific infrastructure and was associated with taxonomic revisions, socioeconomic interest, and disparities in political stability and colonial history. Appropriation of foreign holotypes was greatest in countries with high research investment, strong environmental governance, and historical geopolitical influence.

**Main conclusions:** Global patterns of holotype circulation reflect a persistent geography of scientific inequality. The distribution of taxonomic reference material emerges from the interaction of retention, extraction, and appropriation processes, linking local biodiversity discovery to uneven global scientific capacity. Reducing these inequalities will require investments in taxonomic expertise, institutional infrastructure, and governance frameworks that promote more equitable stewardship of biodiversity knowledge and its material foundations.

## Introduction

A major challenge in contemporary biology is the incomplete documentation of Earth’s biodiversity, as a substantial fraction of species remains unknown to science. This limitation has been called the Linnean shortfall and constrains our ability to characterize eco-evolutionary patterns and to inform conservation planning (Costello et al., 2013; Hortal et al., 2015; Mora et al., 2011). Because species often constitute the operational units of macroecology, biogeography, threat assessments, and environmental governance, undescribed taxa are often invisible to conservation planning and biodiversity policy (Tedesco et al., 2014; Whittaker et al., 2005). Yet species descriptions do more than add names to lists. Each description produces a permanent reference point: the holotype, the name-bearing specimen that anchors the scientific name (Dayrat, 2005). Holotypes are not merely nomenclatural formalities; they are the physical benchmarks against which future taxonomic hypotheses, synonymies, conservation reassessments, and comparative biodiversity studies are validated (International Commission on Zoological Nomenclature, 1999; L. Meyer et al., 2026). Therefore, easy access to them can directly shape the speed, stability, and reproducibility of biodiversity documentation.

The scientific value of holotypes usually depends on physical accessibility. Although digitization, imaging, and molecular tools have expanded remote access to biodiversity information (Hebert & Gregory, 2005; Page et al., 2015), the examination of original material remains crucial for many organisms, especially when diagnostic characters involve fine morphology, osteology, or reproductive structures (Dayrat, 2005; Sangster & Luksenburg, 2015). When holotypes are difficult to access, taxonomic revisions can slow down, misidentifications persist, and undescribed species already housed in collections can remain hidden for years (Fontaine et al., 2012; Goodwin et al., 2015; Guedes et al., 2020). Reptiles make this problem particularly timely. They are the most diverse extant tetrapod group, now exceeding 12,000 described species, and are expected to account for half of all future discoveries of terrestrial vertebrates (Moura & Jetz, 2021; Uetz et al., 2026). Thus, barriers to holotype access can quickly cascade into broader limitations for systematics and conservation.

Emerging evidence shows that access to name-bearing specimens is uneven worldwide, and that holotype extraction is not just a historical legacy but an ongoing feature of modern biodiversity research (Moura et al., 2026; Nakamura et al., 2025; Raja et al., 2021). Countries where new species are most frequently discovered, typically tropical and megadiverse nations, are often not the same countries where their holotypes are ultimately deposited (Nakamura et al., 2025; Raja et al., 2021). This creates a spatial disconnect between biodiversity discovery and the infrastructure needed to validate, revisit, and refine that knowledge over time. The problem is not simply logistical, reflecting a broader geography of scientific power, in which biodiverse regions disproportionately supply specimens and field expertise, while wealthier countries retain museums, collections infrastructure, and long-term control over reference material (Nakamura et al., 2023; Raja et al., 2021). These colonial dynamics in biodiversity science can manifest through multiple pathways, such as authorship exclusion, unequal access to analytical tools, and asymmetrical collaborations (Abreu et al., 2025; Moura et al., 2026; Nakamura et al., 2025). However, the displacement of holotypes may be one of its most durable expressions because it relocates the physical anchor of species identity itself.

Most previous efforts to quantify these asymmetries have primarily relied on aggregate country-level descriptors, such as proportions of retained holotypes, bilateral specimen-flow networks, or national-level indicators of extraction and appropriation (Moura et al., 2026; Nakamura et al., 2025; Raja et al., 2021). These approaches have been instrumental in revealing the broad geopolitical structure of biodiversity knowledge.

However, as widely recognized in ecology, the drivers and patterns of biodiversity are inherently scale-dependent (Levin, 1992), and relationships that appear coherent at aggregate levels may conceal substantial heterogeneity at finer resolutions (McGill, 2010). In this context, country-level summaries may obscure the biological and taxonomic processes that ultimately shapes whether a given holotype stays or leaves. Attributes operating at the level of species (e.g., clade identity, body size, authority size) and type-localities (e.g., on-ground accessibility, endemism, latitude), as well as relational properties between origin and destination countries, may introduce systematic variation that is not captured by aggregated metrics. For example, the same biases that skew sampling toward larger and charismatic taxa (Nekola et al., 2019) may also influence holotype fate. Integrating these complementary levels of inference is key to linking global patterns to local processes, showing that while biodiversity knowledge imbalances are global in scope, they can be locally generated, and therefore require locally grounded solutions.

Here, we analyse global patterns of holotype circulation using a multi-level framework that integrates three complementary units of analysis: individual species and the movement of their holotypes between origin and deposition countries, pairwise connections between countries linked by holotype transfers, and aggregate country-level patterns. We first reconstruct the historical geography of reptile holotype redistribution from 1758 to 2024, characterizing long-term shifts in global retention, concentration, and international specimen flows. We then focus on the contemporary period (1990–2024) to assess how biological, institutional, and geopolitical factors shape whether holotypes are retained locally or deposited abroad (Box 1). By integrating historical and contemporary perspectives, our goal is to understand how colonial legacies continue to structure biodiversity knowledge production and how reducing these asymmetries may contribute to more equitable and efficient species documentation.

#### Box 1. Hypotheses on the drivers of contemporary holotype specimen circulation.

We formulate predictor-specific expectations spanning species-level attributes and country-level conditions, while distinguishing processes operating at origin, destination, and relational levels (i.e., disparity or difference between origin and destination characteristics). Importantly, holotype **retention** and **extraction** are evaluated at both species and country levels, where they represent related but distinct concepts. At the species level, retention refers to whether a holotype remains within its region of origin, whereas extraction refers to the transfer of a holotype to a foreign destination. At the country level, retention corresponds to the proportion of native species whose holotypes are housed domestically, while extraction reflects the proportion of non-native holotypes held in a country’s collections. Because these metrics capture different dimensions of specimen circulation, they may reflect distinct but interconnected processes operating simultaneously within a push-pull dynamic, whereby the same underlying drivers can simultaneously promote both the retention and extraction of specimens.

**Species-level covariates:**

- ***Absolute latitude*** - Tropical regions concentrate a disproportionately larger share of global biodiversity and undescribed taxa relative to temperate regions (Moura & Jetz, 2021; Roll et al., 2017). This imbalance is expected to increase reliance on external expertise, promoting holotype extraction from low-latitude regions. Conversely, where local scientific capacity is stronger, this same biodiversity concentration may enhance holotype retention. Overall, we predict that holotype extraction will be disproportionately associated with lower latitudes, whereas retention will depend on local capacity mediating this effect.
- ***Body size* -** Larger-bodied species tend to be more detectable and garner more scientific attention (Guedes et al., 2023; Moura & Jetz, 2021), which may increase international research attention and holotype extraction. However, larger specimens may also impose greater transport and curation costs, potentially favouring local retention. We therefore expect body size to influence holotype circulation, but the direction and magnitude of this relationship remain an open empirical question.
- ***Collector inclusion*** - The involvement of field collectors as co-authors may reflect stronger local engagement (Abreu et al., 2025; Guedes et al., 2020), and is expected to increase the likelihood that type material is curated within the origin region. We expect higher holotype retention and lower extraction when at least one collector is included among the authors of the formal species description.
- ***Collection site protection status*** - Protected areas are generally associated with formal governance structure, permitting requirements, and institutional oversight of biological sampling activities (e.g., SNUC, 2000). We expect holotypes collected within protected areas to exhibit greater retention within the country of origin, reflecting reduced opportunities for unregulated specimen removal.
- ***Number of authors*** - Larger and more diverse collaborations increase the probability of including domestic researchers (Joppa et al., 2011), reducing exclusionary practices (extraction) and favouring deposition (retention) in origin countries.
- ***Socioeconomic interest*** - Species that are culturally salient, useful, or charismatic attract disproportionate scientific attention (Mammola et al., 2023; Troudet et al., 2017), increasing the likelihood of international study and holotype extraction. Alternatively, holotype retention may also increase if specimen socioeconomic value is properly recognized locally (Moura et al., 2026).
- ***Taxonomic revision*** - During revisionary work, taxonomists may visit collections and borrow specimens to apply multiple diagnostic tools and techniques (Guedes et al., 2024; Moroti et al., 2026). As a result, specimens may be permanently transferred to well-equipped institutions, increasing the probability of holotype deposition outside the origin region.
- ***Year of description*** - Contemporary taxonomy increasingly reflects more inclusive and capacity-building practices, contrasting with historically extractive patterns (Guedes et al., 2024; Nakamura et al., 2025). We expect earlier-described species to show higher holotype extraction and lower retention.

**Country-level covariates:** Country attributes may influence holotype circulation when considered at the level of origins or destinations, but their effects may also be shaped by disparities between countries. Such imbalances are expected to reduce local retention capacity while increasing the extractive potential of more advantaged nations.

- ***Colonial history*** - Colonial legacies shape structural pathways of specimen movement, including established institutional ties, legal norms, and authority over (Moura et al., 2026; Nakamura et al., 2025; Raja et al., 2021). We expect former colonial powers to retain a greater proportion of their own holotypes while also accumulating imported holotypes originating from other countries. In contrast, former colonies are expected to exhibit lower retention and higher export of locally collected holotypes.
- ***Endemism richness*** - Countries with high endemism tend to attract disproportionate international research effort due to their unique biodiversity (C. Meyer et al., 2015, 2016). This primarily acts as a biological attractor of research effort, increasing extraction risk independently of local capacity. We investigated whether local systems can convert this attention into sustained local taxonomic activity.
- ***Environmental governance*** - Strong environmental regulation reflects a country’s ability to protect biodiversity, enforce collection permits, and promote local stewardship of biological resources (Moura et al., 2026), reducing opportunities for holotype extraction and limiting unregulated export (Cisneros et al., 2022). We expect higher holotype retention under stronger biodiversity and environmental governance. Conversely, weaker governance (or disparities in governance between countries) may facilitate extraction by reducing oversight and allowing external actors to access and relocate specimens more easily.
- ***Human capital*** - Availability of trained specialists increases national capacity to perform biodiversity research (dos Santos et al., 2020), and thus describe and curate biodiversity locally (Moura et al., 2026). We expect countries with higher human capital (or researcher per capita) to show higher holotype retention. In contrast, holotype extraction is expected to reflect disparities in human capital between countries, with specimens more likely to move from regions with lower taxonomic capacity to those with greater expertise.
- ***Institutional availability*** - Availability of biodiversity institutions, reflected either by the number of institutions or by their collection size (i.e., the number of specimens they house), provides the physical and logistical infrastructure necessary for specimen deposition and long-term curation (Guedes et al., 2023; Moura et al., 2026). We expect greater institutional availability to increase holotype retention and facilitate the accumulation of foreign type material, thereby increasing specimen extraction.
- ***Research investment*** - Financial capacity to sustain scientific activity influences the scale and continuity of biodiversity research (Guedes et al., 2023; C. Meyer et al., 2015). We expect higher investment to increase a country’s ability to retain locally generated holotypes, while also enhancing its capacity to acquire type material through international collaboration.
- ***Secure conditions*** - Political stability allows for high levels of public safety and lack of armed conflicts, creating favourable conditions for biodiversity sampling (C. Meyer et al., 2015; Raja et al., 2021). We expect more peaceful countries to exhibit higher holotype retention. In contrast, instability may disrupt local research systems and increase reliance on external collaborators, promoting holotype extraction toward more stable countries where long-term curation is more secure.

## Methods

### Species-level data

We examined 12,314 descriptions of reptile species published from 1758 to 2024, as informed by The Reptile Database (Uetz et al., 2026). We compiled information on holotype countries of origin and deposition primarily from the recently published global catalogue of reptile primary type specimens (Uetz et al., 2019), supplemented by recent data mobilization efforts (Guedes et al., 2020, 2024) and targeted searches of original descriptions, taxonomic revisions, and museum records for species lacking complete information. For each species holotype, we compiled information on both the country of origin and the country of deposition, including the corresponding ISO alpha-3 country codes following the United Nations geopolitical framework (United Nations Statistics Division, 2024). Overall, we retrieved data for 11,645 reptile holotypes, with 669 species discarded to uncertain information on either origin- or destination-country. Country assignments were based on contemporary political boundaries and geopolitical classifications following *World Administrative Boundaries* (2025). Consequently, historical colonial territories and geopolitical transitions were represented according to present-day political units rather than the political entities existing at the time of collection or deposition.

Countries were assigned to one of eight geopolitical regions: (i) Sub-Saharan Africa; (ii) Latin America and the Caribbean; (iii) Northern America; (iv) Australia and New Zealand; (v) Central, Eastern, and Southern Asia; (vi) Europe; (vii) Near East and Northern Africa; and (viii) Southeast Asia and Pacific Islands. To improve geographic resolution, the Russian Federation was treated separately from Europe. We followed Abreu et al. (2025) and classified as Global North those countries located in Northern America, Europe, Australia and New Zealand, and the Russian Federation, whereas all remaining regions were classified as Global South, except for Japan, South Korea, Israel, and French Guiana, which were treated as part of the Global North despite being geographically embedded within predominantly Global South regions.

To characterize long-term changes in reptile holotype redistribution, we quantified the proportion of holotypes exported or retained per decade between 1758 and 2024. We additionally aggregated species records into directional origin–destination flows among geopolitical regions across six historical periods (1750–1799, 1800–1849, 1850–1899, 1900–1949, 1950–1989, and 1990–2024) to reconstruct major shifts in global holotype redistribution through time. Preliminary inspection of these temporal trends indicated substantial changes in retention and redistribution dynamics after the late twentieth century, supporting the use of the 1990–2024 interval as representative of contemporary taxonomic and socioeconomic conditions, in contrast to the expedition-based and colonial scientific structures that predominated during the nineteenth and early twentieth centuries. Operations were conducted in R software v. 4.5.0 (R Core Team, 2025).

For species described between 1990 and 2024, we compiled detailed metadata from species description. Specifically, we obtained type-locality’s latitude and longitude. When only textual locality descriptions were available (e.g., protected areas or municipalities), approximate coordinates were assigned using gazetteers or Google Earth (https://earth.google.com). We extracted four continuous variables: (1) year of description; (2) number of authors; (3) absolute latitude of type-locality. We also extracted (4) maximum body mass for each species, using recently published datasets (Feldman et al., 2016; Guedes et al., 2024; Moura et al., 2024) and original descriptions. Although these body size values were not always derived directly from holotype specimens, they should still provide a reliable approximation of broad-scale patterns because intraspecific variation is generally expected to be smaller than interspecific variation.

We also defined four binary variables (1 = yes, 0 = no) informing: (5) whether the description was based on a taxonomic revision, considered only when the study explicitly used the term “revision” (or similar) in the title, abstract, keywords, or main text; (6) whether collectors of holotype specimen were included as co-authors, used as a proxy for inclusivity toward field personnel; (7) whether the holotype was collected within an protected area, as defined by World Database on Protected Areas (UNEP-WCMC & IUCN, 2026). To refine this classification temporally, we retrieved the year of establishment associated with each protected area and considered the type-locality as protected only when the corresponding protected area had already been established at the time of holotype collection. Finally, we recorded (8) whether the species exhibits socioeconomic interest. To classify socioeconomic interest, we used genus-level evidence of biological exploitation based on the IUCN Threats Classification Scheme v. 3.3 (IUCN - International Union for Conservation of Nature, 2026). Specifically, we used the *rredlist* (Scott Chamberlain, 2020) to extract data on threat type 5 (“Biological Resource Use”), treated at the species-level as a binary indicator, and then aggregated these values at the genus-level as the proportion of species affected. Genera in which more than 50% of species were associated with this particular threat were classified as having socioeconomic interest.

### Country-level data

For each country, we extracted nine variables reflecting research capacity, biodiversity context, and historical and governance conditions. Specifically, we quantified: (1) the number of biodiversity institutions (e.g., museums, herbaria, universities) using spatial data from the *CoordinateCleaner* package (Zizka et al., 2019), complemented with manual curation from Open Alex database (https://api.openalex.org/) for countries lacking coverage. (2) The number of preserved reptile specimens collected per country, derived from 2,136,273 occurrences after removing records with no coordinates or duplicated entries with the same catalogue number and repository institution (GBIF, 2026). (3) Endemism richness, calculated as the country-level average of grid-cell values derived from mapping species ranges across a global 110×110 km cylindrical equal-area grid. For each cell, endemism richness was computed as the sum of the inverse range sizes of all species whose distribution maps overlapped that cell (Kier & Barthlott, 2001), based on spatial data from TetrapodTraits v. 2.0.1 (Moura et al., 2024).

We used the *wb_data* function from the *wbstats* R package (Piburn, 2020) to retrieve World Bank Open Data indicators on (4) research and development expenditure as a percentage of GDP and (5) researchers per million inhabitants (hereafter, researchers per capita), both averaged across the period 1990–2024. Other four indications were extracted from the Quality of Government (QoG) database (Teorell et al., 2025), namely: (6) Human Capital Index, evaluating how well countries develop education and health for productivity, with higher values indicating greater human capital development; (7) Biodiversity and Habitat Issue category, used as a national indicator of environmental governance, with higher values indicating stronger biodiversity conservation and protected-area performance; (8) Global Peace Index, measuring national peacefulness based on conflict, safety, and militarization, with higher values indicating lower levels of peace and stability; and (9) Colonial history. The latter is a binary variable indicating whether a country has experienced a history of external colonial rule (former colony = 1) or not (0). Each country’s value was averaged across the available QoG time series from 1990 to 2024, except for colonial history, which is time invariant.

Because country-level socioeconomic data are unevenly available (particularly for Global South nations), we avoided discarding countries with missing values by implementing a structured imputation procedure. Missing values for continuous predictors were imputed using median values calculated within relevant geopolitical groupings, prioritizing Least Developed Countries (LDCs) when applicable, and otherwise using macroregional medians (Caribbean, South America, Micronesia, Melanesia, Polynesia, Eastern Africa, Northern Africa, Southeast Asia, Central Asia, Western Africa, Middle Africa, Western Asia, Eastern Asia). This procedure preserved the full set of countries while minimizing bias associated with systematic data gaps. All imputation steps were recorded to allow post hoc quantification of the number of countries and variables affected.

### Metrics of holotype movement dynamics

We quantified holotype movement at three complementary levels: species-level retention status, bilateral flows between country pairs, and aggregate country-level metrics. At the species-level, we defined holotype retention as a binary variable based on whether the country of the type-locality matched the country of the housing collection (1 = retained, 0 = exported). At the bilateral-level, we aggregated species-level records into origin-destination pairs, counting the number of holotypes collected in each origin country and housed in each destination country to construct a weighted network of directional flows across country pairs.

At the country-level, we quantified holotype retention as a binomial outcome based on counts of successful versus unsuccessful retention events. Specifically, for each country, we defined the number of retained holotypes as those whose type-locality and housing country matched, and the number of exported holotypes as those housed abroad. These counts were jointly modelled using a beta-binomial framework (see Data Analysis). Holotype appropriation was modelled as the number of foreign holotypes housed, with an offset term corresponding to the logarithm of the total number of species housed to capture the relative accumulation of foreign holotypes within a country.

In addition, we quantified countries’ positions within the global holotype exchange network using centrality-based metrics. We built a bipartite network linking countries of type-locality to countries hosting holotype collections, with edge weights corresponding to the number of specimens exchanged between each pair of countries. At the node level, we estimated Katz centrality to quantify each country’s structural influence within the network, accounting for both direct connections and indirect pathways through other countries (Katz, 1953). The attenuation parameter (α = 0.1) was used to down-weight contributions from longer paths (range: 0–0.2), and calculations were implemented using the *katzcent* function in the *centiserve* R package (Jalili et al., 2015). Exploratory analyses indicated that inflow and outflow metrics were not influenced by country area (See Data Availability).

### Data Analysis

To improve comparability among predictors on different scales, variables describing research capacity and biodiversity context (e.g., number of collected specimens, number of research institutions, number of researchers per capita, research expenditure, and endemism richness) were log_10_-transformed to reduce skewness. All continuous variables were subsequently centred and standardized (mean = 0, SD = 1) to allow direct comparison of effect sizes. Multicollinearity among predictors was assessed using Variance Inflation Factors (VIF) implemented in the *performance* package (Lüdecke et al., 2021), and variables with high collinearity were excluded if VIF > 10 (Kutner et al., 2004).

At the species-level, we modelled the probability that a holotype is retained in its country of origin using a binomial generalized linear mixed-model (GLMMs) (Zuur et al., 2009) with logit link implemented in the *glmmTMB* package (Brooks et al., 2017). Fixed effects included species-level attributes (year of description, number of authors, taxonomic revision, collector authorship, collection site protection status, absolute latitude, and socioeconomic interest), as well as country-level indicators describing research capacity (number of collected specimens, number of research institutions, researchers per capita, research spending as % of GDP), biodiversity context (endemism richness and Biodiversity and Habitat Issue), and governance and historical factors (Global Peace Index, Human Capital Index, and colonial origin). Country of origin was included as a random effect to account for non-independence among species described from the same country.

At the bilateral level, we modelled directional holotype flows between origin–destination country pairs using GLMMs with a truncated negative binomial distribution. Domestic flows (origin = destination) were excluded to avoid structural tautologies. We fitted three complementary models: (i) an origin-country model evaluating how attributes of source countries influence outward holotype flows, with origin country included as a random effect; (ii) a destination-country model assessing how attributes of destination countries influence incoming flows, with destination country included as a random effect; and (iii) a disparity model testing how differences between origin and destination countries (Δ = destination value – origin value) affect the intensity of holotype exchange, with origin country included as a random effect. Separating these models avoids multicollinearity and circularity that would arise from jointly including origin, destination, and difference-based predictors.

At the country-level, we modelled aggregate patterns of holotype retention, appropriation, and network centrality. Only countries with at least three species described or three deposited holotypes were included in these analyses. Retention was modelled as the number of retained versus exported holotypes using a beta-binomial distribution to account for overdispersion in the binomial response (Harrison, 2015). Importantly, retention was estimated conditional on the total number of holotypes associated with a country, rather than as a simple percentage, allowing countries with similar proportional retention but different magnitudes of taxonomic activity to be distinguished. Holotype appropriation (number of imported holotypes) was modelled using a negative binomial distribution, with a log-transformed offset term for the total number of housed species, thereby modelling the relative accumulation of foreign holotypes while accounting for collection size (Hardin & Hilbe, 2018). Finally, inflow and outflow centrality were modelled using gamma distributions with log link functions (Zuur et al., 2009). Because inflow centrality exhibited substantial right-skewness, we log-transformed the response variable prior to analysis to improve model fit before fitting a gamma model with a log link.

Model fit was evaluated using R² metrics suited to each model structure and error distribution. For the GLMMs at the level of either species or country-pairs, we used Nakagawa R² to inform explanatory power of fixed effects only or fixed + random effects. For the beta-binomial retention model at the country-level, we used McFadden’s R². The explanatory power of all other country-level models (negative binomial model of holotype appropriation, Gamma models of inflow and outflow centrality) were assessed via Nagelkerke’s R². The R² metrics were computed using the *performance* package (Lüdecke et al., 2021). Model assumptions were checked using simulation-based residual diagnostics implemented in the *DHARMa* package (Hartig, 2022).

## Results

We traced origin and destination of holotypes for nearly 95% of reptile species worldwide. Overall, these holotypes were collected across 176 countries and housed in 76 nations. The Global North harboured 13.9% (N = 1619) of species described and 76.4% of reptile holotypes (N = 8893), whereas 86.1% of all species were discovered in the Global South (N = 10,026), which housed 23.6% (N = 2752) of holotypes.

### Historical pattern of holotype circulation

Across most of the historical record, holotype-destination countries consistently exhibited lower retention of holotypes (Fig. 1a). Only in the 1990s that proportion of retained holotypes outnumbered exported holotypes. Overall, historical patterns of reptile holotype redistribution revealed persistent transfers from biodiversity-rich regions in the Global South toward repositories located predominantly in Europe during the eighteenth and nineteenth centuries, later expanding to Northern America during the late nineteenth and early twentieth centuries, and more recently including Australia and New Zealand after the 1950s (Fig. 1b). Across all decades until the 1980s, Global North nations housed at least 80% of reptile holotypes worldwide, although this proportion declined to 64% in the 1990s and continued decreasing thereafter. Across major reptilian clades, crocodilians showed the highest exportation rate (96.2% of all holotypes), followed by Dibamoidea (92.9%), snakes (77.9%), and chelonians (77.8%), whereas the lowest exportation rate was observed in Lacertoidea lizards (58.5%) and Gekkota (62.9%, Fig. S1). Although all major reptilian clades showed higher exportation than retention of holotypes, but exportation rates have declined over time.

**Figure 1.**
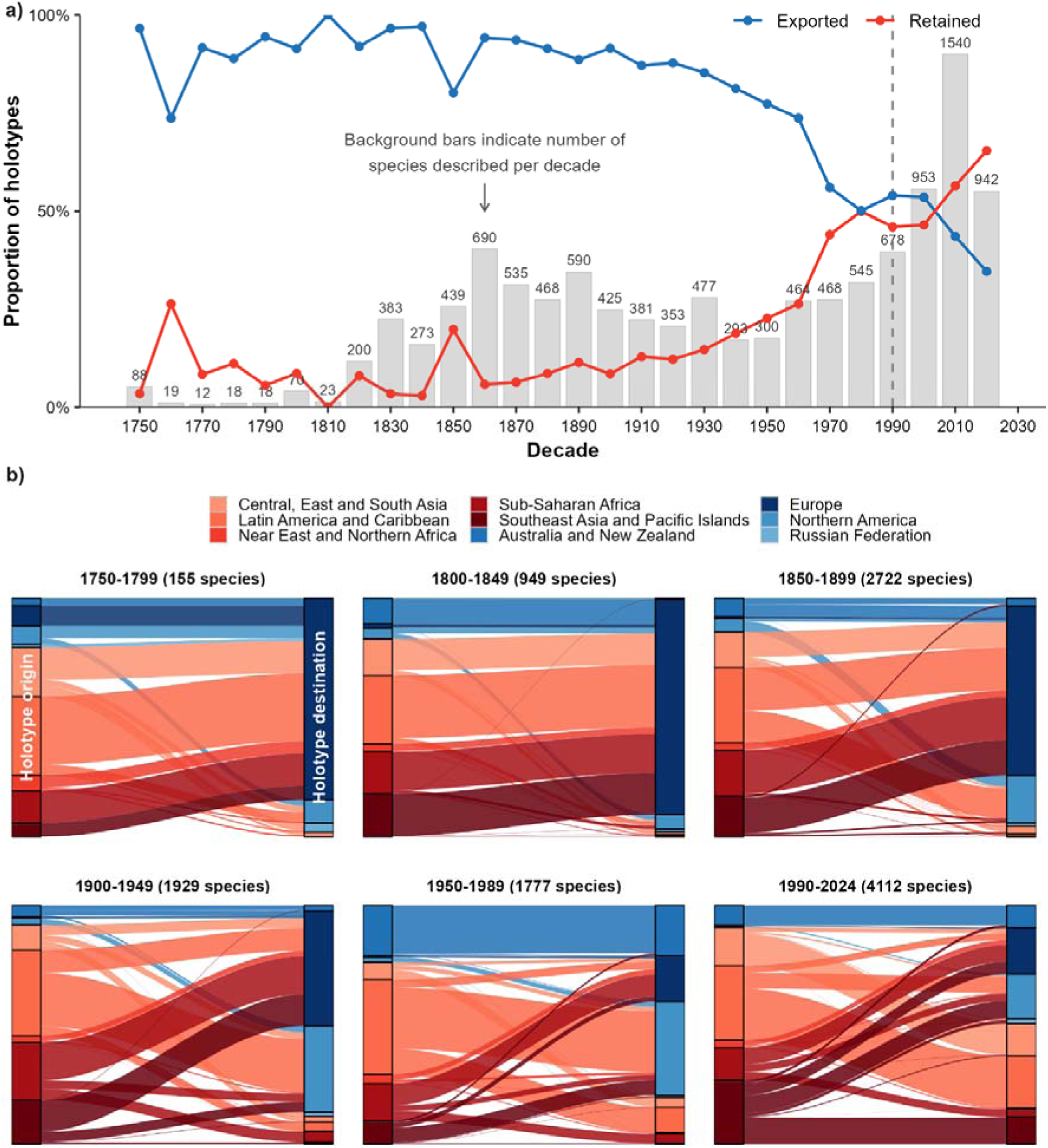
Historical dynamics of global reptile holotype flows and distributional equity among countries. (a) Temporal trends in the proportion of reptile holotypes retained or exported from the 1750s to the 2020s. Grey background bars represent the total number of reptile holotypes described per decade, illustrating the strong post-1990 acceleration in species descriptions. (b) Sankey diagrams depicting the global circulation of reptile holotypes among major geopolitical regions across six historical periods. Flows connect the geopolitical regions of origin (left) and destination (right) of species holotypes, with ribbon widths proportional to the number of holotypes transferred.

### Contemporary pattern of holotype circulation

Focusing on the 1990–2024 period, 90.4% (N = 3716) of all reptile species were described from the Global South, which housed 49.9% of holotypes (N = 2054). The Global North harboured 9.6% (N = 396) of species described and 50.1% of reptile holotypes (N = 2058). Species described spanned all major geopolitical regions, with Latin America and the Caribbean accounting for the largest share at 31.3% (1287 species), followed by Southeast Asia and Pacific Islands (26.8%; 1102), Central, East and South Asia (16%; 658). Reptile holotypes were deposited mostly in Latin America and the Caribbean 21.7% (892 species), Europe (19.4%; 796), and Northern America (18.7%; 770). Nevertheless, 81.3% of holotypes housed in the Global North collections were extracted abroad, contrasting with the 9.6% of non-native holotypes deposited in the Global South collections (Fig. 2).

**Figure 2.**
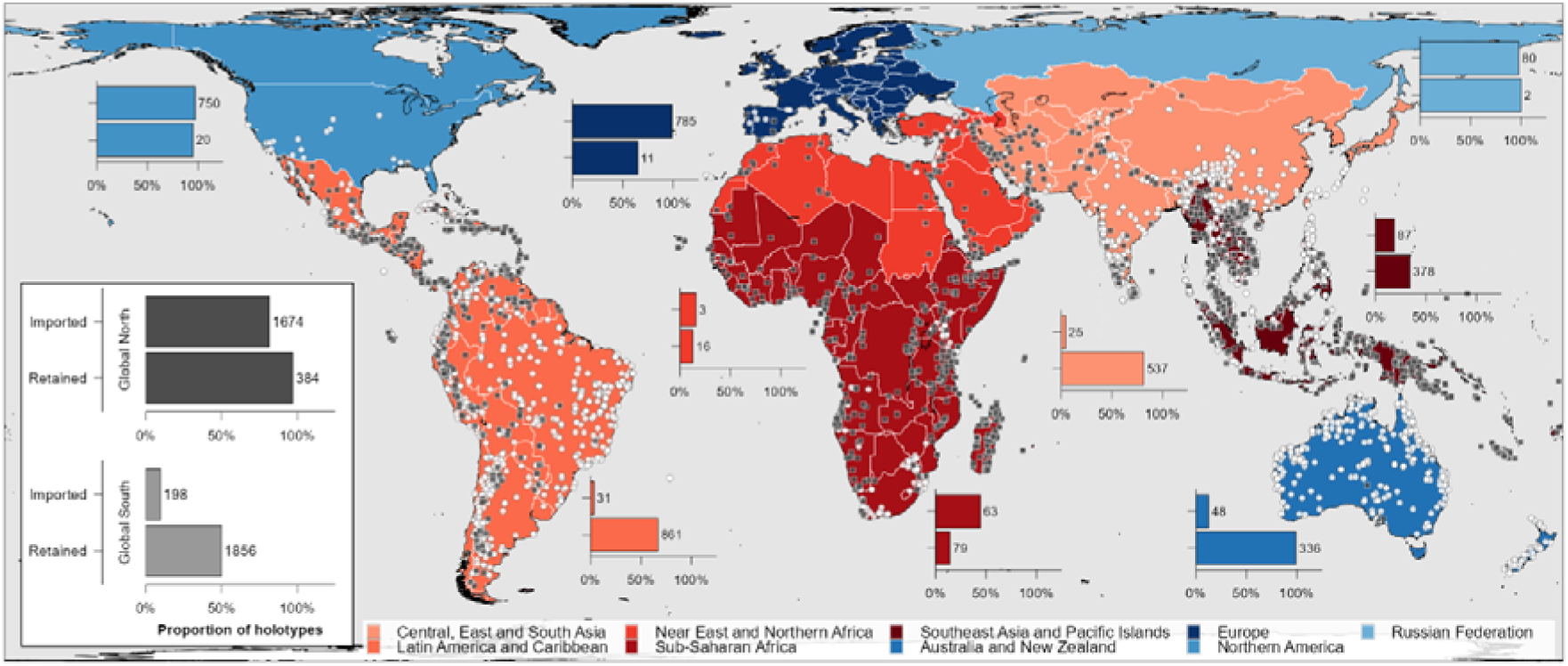
Proportion of reptile holotype specimens retained or imported across geopolitical regions. Points indicate type-localities of reptile species described between 1990 and 2024 whose holotypes were retained (white circles) or exported (grey squares). Barplots show the proportion of holotypes that were retained or imported in each macropolitical region. Numbers adjacent to bars are holotype counts. Reddish and bluish colours distinguish geopolitical regions associated with the Global South and Global North, respectively. The map uses an equal-area projection.

Of the 146 holotype origin-countries, 139 had at least one holotype housed abroad, and 92 experienced complete extraction (i.e., 100% of holotypes exported), with 90 of those located in the Global South (the two exceptions were Montenegro and Slovenia, each represented by a single species whose holotype was deposited in another European country). Network analyses further reveal a strongly asymmetric flow structure, with holotypes predominantly moving from biodiversity-rich regions in the Global South to institutions in the Global North (Figs 3-4). Germany, United States, France, and the United Kingdom emerged as central hubs of holotype inflow and accumulation, whereas key source regions include Madagascar and Southeast Asian nations such as Indonesia, Thailand, and Vietnam, with additional contributions from a few Latin America countries (Mexico, Panama, Peru, and Honduras). Countries with the highest holotype outflow (Katz out-centrality) are exclusively in the Global South, while those with the highest holotype inflow (Katz in-centrality) are overwhelmingly in the Global North (Figs 3-4 and S2-S5).

**Figure 3.**
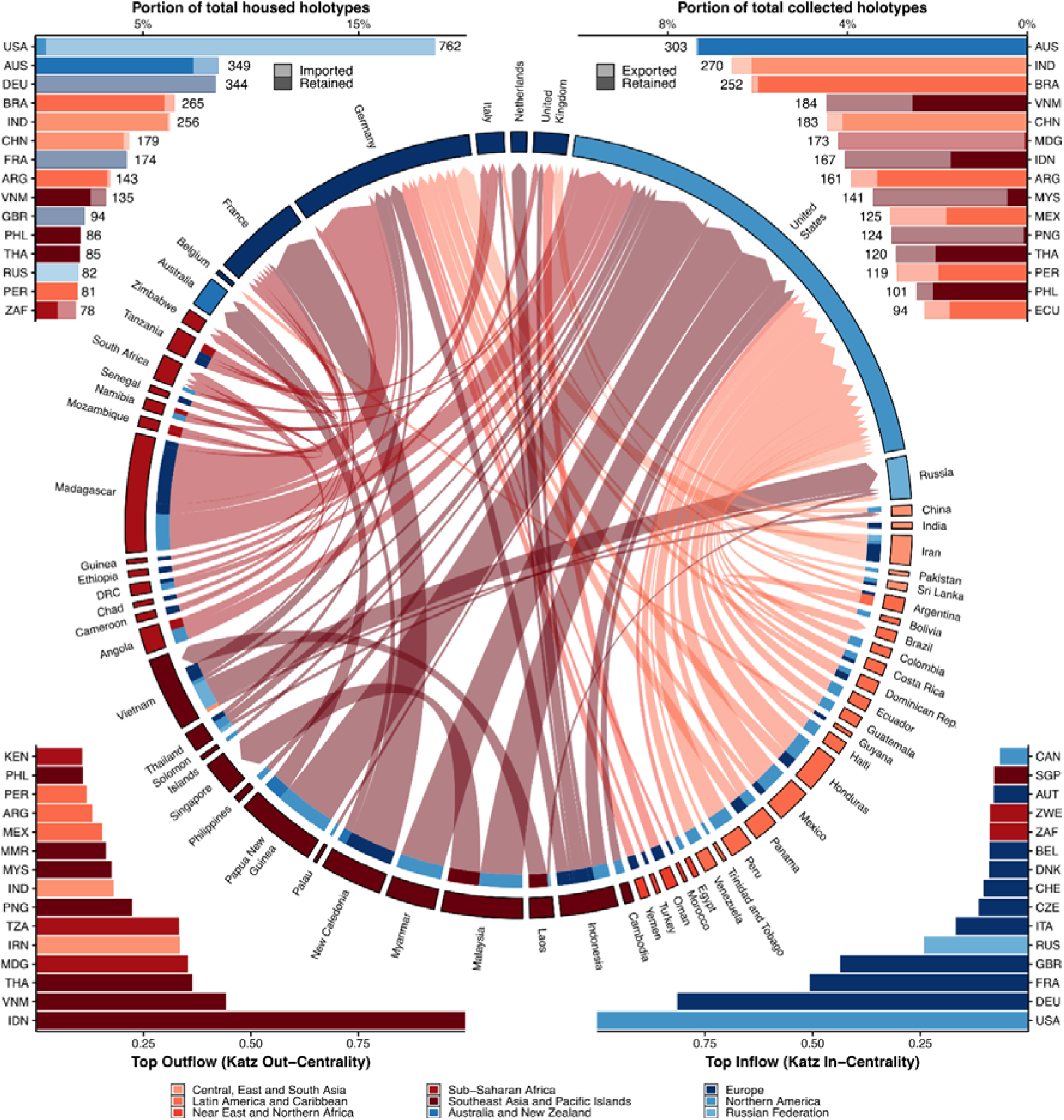
Global outflow of reptile holotype specimens across countries (1990-2024). Lateral barplots show top-15 countries (ISO-alpha3 code) in holotype destination (top-left), origin (top-right), outflow (bottom-left), and inflow (bottom-right). High inflow centrality indicates countries exerting extractive influence in parachute science networks, while high outflow centrality reflects greatly exploited countries. For better visualization only countries with at least five holotypes are shown. Arrows represent origin-to-destination pathways of holotypes between countries (self-links omitted). Reddish and bluish colours distinguish geopolitical regions associated with the Global South and Global North, respectively. See Figs S2-S3 for complete country-rankings and Fig. S4 for a representation of including holotype retentions.

**Figure 4.**
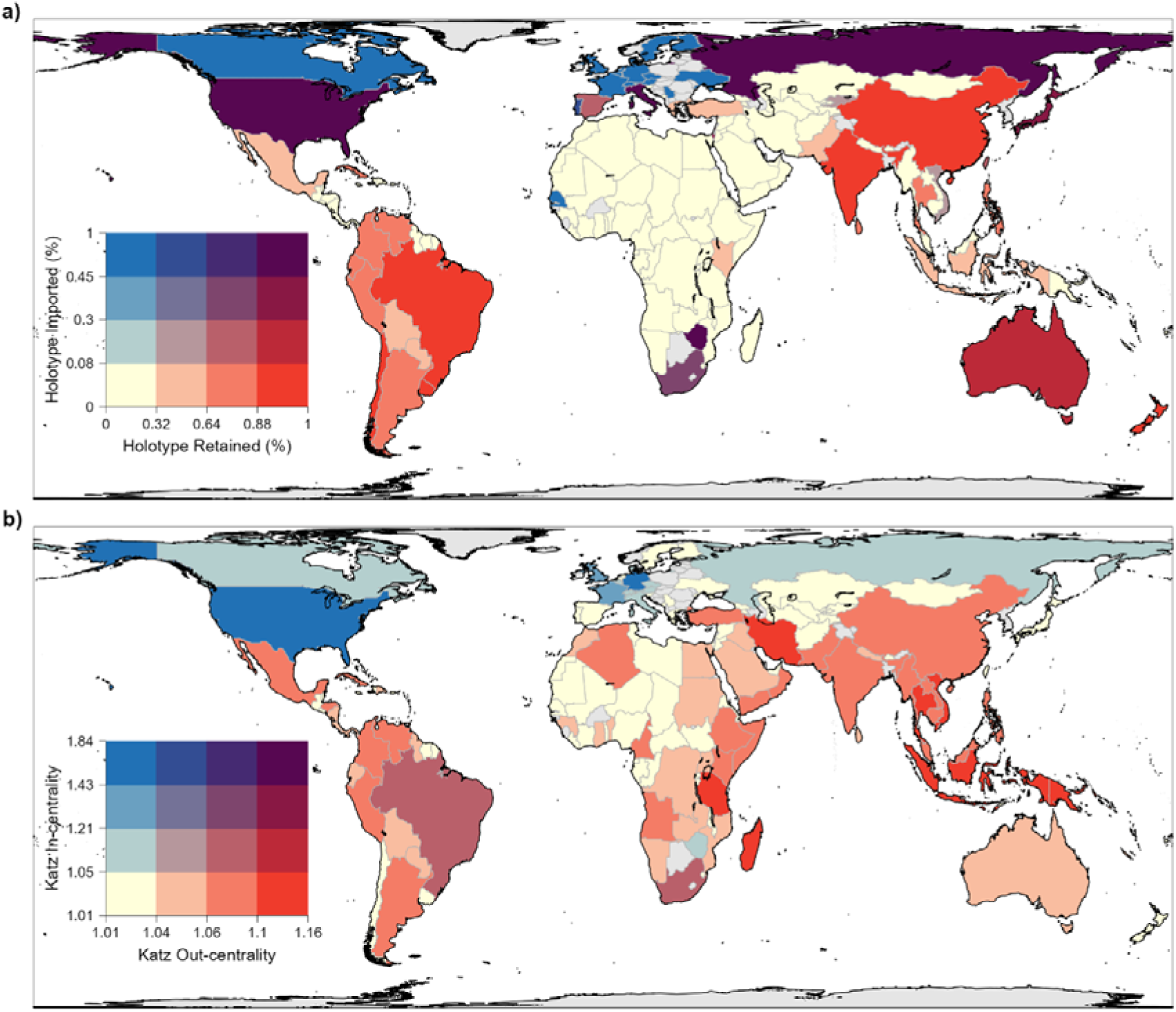
Geographical patterns of reptile holotype specimens transfer across countries (1990-2024). a) Per country proportion of holotypes imported or locally retained. b) Holotype flows across countries as indicated by Katz In-centrality and Katz Out-centrality. Colours in the bivariate maps indicate the combined values of the two variables, as shown in the inset legend. Countries lacking information for one of the two dimensions were assigned the lowest value for visualization purposes only. Countries with no holotypes discovered or housed are shown in grey. Maps were drawn using an equal-area projection. Individual spatial patterns for each metric are shown in Fig. S5.

### Determinants of holotype circulation patterns

Variation in holotype retention was well explained by the model, with fixed effects accounting for 64.1% of the variance (marginal R²), increasing to 82.2% (conditional R²) when including variation among origin countries (Fig. 5a, Table S3). Including collectors as co-authors had one of the strongest positive effects, increasing the odds of local retention by ∼180% (OR ≈ 2.80, p < 0.001, Fig. 5d). More recent species descriptions, collection sites overlapping protected areas, larger author teams, and species with socioeconomic interest were also associated with greater retention odds by approximately 85%, 42%, 27%, 36%, respectively). In contrast, species described through taxonomic revisions and larger bodied were less likely to be retained locally, reducing retention odds by approximately 50% and 13%, respectively (all p < 0.05, Table S3). Other species-level predictors showed weak or non-significant effects. Among country-level predictors, human capital and the number of institutions showed particularly strong positive effects, increasing retention odds by approximately 366% and 1,313%, respectively (both p < 0.001, Table S3).

**Figure 5.**
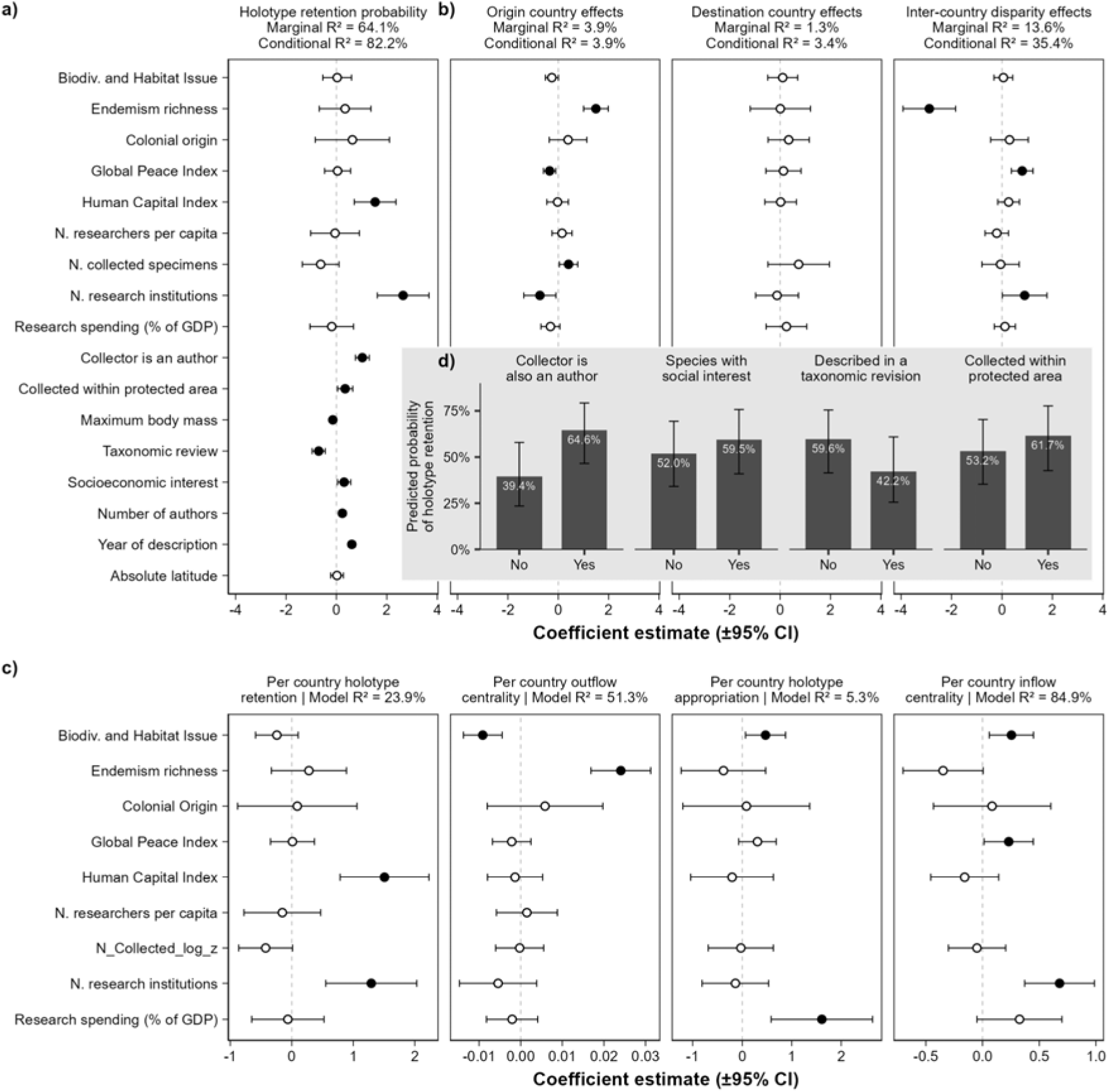
Multilevel determinants of holotype circulation dynamics (1990-2024). (a) Fixed-effect estimates from the species-level mixed model of holotype retention. (b) Fixed-effect estimates from mixed models evaluating origin-country, destination-country, and country-disparity effects (Δ = destination value – origin value) on bilateral holotype exportation flows. (c) Fixed-effect estimates from country-level models of holotype retention, outflow centrality (the degree of holotype extraction by different foreign institutions), foreign holotype appropriation, and inflow centrality (the country’s extractive influence). In panels a-c, filled circles indicating significant effects (p < 0.05). (d) Model-adjusted probabilities of holotype retention across binary species-level attributes, estimated from the species-level GLMM while holding all remaining predictors constant. See tables S3-S6 for additional details on model diagnostics. Researcher per capita was omitted from some models due to high multicollinearity.

The GLMMs of holotype transfers between country pairs showed limited explanatory power when considering attributes of either the origin countries (fixed-effects R² = 3.9%) or the destination countries (R² = 1.3%) separately. In contrast, bilateral holotype flows were better explained by intercountry differences in these attributes (Δ = destination value − origin value), which increased explanatory power to R² = 13.6% (Fig. 5b, Table S4). Exports were also more frequent from countries with larger numbers of collected specimens, but less frequent from countries with more research institutions. Among disparity-based predictors, larger differences in political stability (std. model coefficient for Global Peace Index β = 0.80 ± 0.22, p < 0.001) and research infrastructure, particularly the number of research institutions (β = 0.90 ± 0.45, p = 0.047), increased bilateral holotype flows, whereas larger disparities in endemic richness reduced exports. Overall, bilateral flows were structured mostly by intercountry disparities and origin-country conditions than by attributes of destination countries.

Country-level models showed contrasting explanatory power across response variables, with inflow and outflow centrality being relatively well explained (R² = 84.9% and 51.3%, respectively), whereas country-level holotype retention (R² = 23.9%) and especially appropriation (R² = 5.3%) were less strongly associated with national socioeconomic conditions (Fig. 5c, Tables S5–S6). Consistent with the species-level analyses, the beta-binomial retention model indicated that country-level retention increased with human capital (Human Capital Index; β = 1.51 ± 0.37, p < 0.001) and the number of research institutions (β = 1.29 ± 0.38, p = 0.001). Across the interquartile range of these predictors, countries with higher human capital and greater institutional availability exhibited approximately sevenfold and sixfold higher odds of retaining their own holotypes, respectively.

Environmental governance (Biodiversity and Habitat Issue) emerged as the most consistent predictor of country-level network structure, being associated with three circulation metrics (Fig. 5c, Tables S5-S6). Countries with stronger environmental governance exhibited lower outflow centrality (β = −0.009 ± 0.002, p < 0.001), indicating reduced susceptibility to foreign extraction, but showed greater holotype appropriation (β = 0.47 ± 0.21, p = 0.023) and higher inflow centrality (β = 0.25 ± 0.10, p = 0.010). Beyond these effects, outflow centrality increased with endemism richness (β = 0.024 ± 0.004, p < 0.001), whereas holotype appropriation was greater in countries with higher investment in research (research spending as a percentage of GDP; β = 1.61 ± 0.52, p = 0.002). Similarly, inflow centrality increased with the number of research institutions (β = 0.68 ± 0.16, p < 0.001) and greater political stability (Global Peace Index; β = 0.23 ± 0.11, p = 0.037).

## Discussion

Global biodiversity documentation remains deeply uneven, with species-rich regions often lacking proportional scientific capacity (Faxon & Chapman, 2025; Hughes et al., 2021). More than three quarters of all reptile holotypes are housed in Global North collections despite most new species being discovered in the Global South, reinforcing the concentration of biodiversity data in wealthier nations (Moura et al., 2026; Nakamura et al., 2025; Park et al., 2023). The geography of holotype distribution emerges from the interplay between retention and extraction, driven by distinct but overlapping biological, institutional, and geopolitical factors operating across scales. Retention is more likely where scientific capacity and institutional infrastructure are stronger, particularly for species described outside revisionary studies and with higher socioeconomic interest. In contrast, extraction is associated with biodiverse regions characterized by high endemism, limited research infrastructure, and secure conditions that facilitate externally led research, with intercountry differences in these variables also playing a major role in holotype extraction. More broadly, stronger environmental governance was consistently associated with reduced holotype outflows and greater participation in international specimen networks.

Our temporal analyses reveal a persistent geographic inequality of reptile holotypes in a relatively small set of repository countries, with Global North collections housing more than 80% of all holotypes until the 1980s (Fig. 1). This pattern mirrors the biodiversity knowledge divide recently documented for freshwater fishes, where name-bearing specimens remain disproportionately concentrated in museums located in wealthier nations (Nakamura et al., 2025). Throughout most of the historical record, destination countries exhibited substantially lower geographic equity than origin countries, indicating that biodiversity was sampled broadly but deposited in relatively few nations. For example, 73 collections house more than 90% of all primary reptile types worldwide and 22 of the 25 largest collections (as measured by holotype counts) are in the Global North (Uetz et al., 2019), highlighting the strong concentration of taxonomic reference material. Despite this long-standing concentration, holotype retention increased after the mid-twentieth century, coinciding with the emergence of new repository hubs in countries such as Brazil, India, China, Vietnam, and Thailand (Fig. S4). This shift may help reduce barriers to accessing name-bearing specimens in the Global South, where more than 90% of future tetrapod discoveries are expected to occur (Moura & Jetz, 2021, 2024).

Species- and country-level analyses indicate that holotype retention is favoured by integration between local fieldwork personnel and scientific capacity and infrastructure (Fig. 5). Stronger scientific infrastructure increases biodiversity documentation through increased research effort (dos Santos et al., 2020; Guedes et al., 2023), sampling intensity (C. Meyer et al., 2015; Oliveira et al., 2016), and inventory coverage (Carvalho et al., 2023). Beyond retaining name-bearing material within origin countries, local participation may also accelerate biodiversity documentation (Moura et al., 2026), as species are described more rapidly when collectors contribute to taxonomic studies, reducing specimen shelf time in collections (Guedes et al., 2020). Indeed, species whose collectors participated as authors were substantially more likely to have their holotypes retained locally (64.6% versus 39.4%; Fig. 5d), and domestic authors were included in 76% of descriptions with retained holotypes but only 10% of those with exported holotypes. Moreover, species of greater socioeconomic interest showed higher probabilities of local retention (59.5% versus 52%; Fig. 5d), indicating that societal preferences known to shape biodiversity knowledge biases (Troudet et al., 2017) also affect where holotypes are ultimately deposited.

Holotype extraction was concentrated in regions with high endemism and limited scientific infrastructure, reinforcing evidence that biodiversity knowledge production remains spatially decoupled from the regions where biodiversity is concentrated, as documented for research effort, species occurrence and inventory data (Guedes et al., 2023; C. Meyer et al., 2015; Titley et al., 2017). Regions with high endemism also tend to harbour substantial hidden or cryptic diversity (e.g., Burriel-Carranza et al., 2025; Xu et al., 2024); particularly in tropical clades where species boundaries remain incompletely resolved and taxonomic uncertainties are greater (Freeman & Pennell, 2021; Guedes et al., 2025). Resolving this cryptic diversity often requires integrative taxonomic approaches (Padial et al., 2010), which depend on costly analytical tools and extensive comparative material, making revisionary research more feasible in scientifically well-resourced institutions (Abreu et al., 2025). Accordingly, species described in revisionary studies were predicted to have lower probabilities of local holotype retention (59.6% versus 42.2%; Fig. 5d), suggesting that revisionary workflows may intensify transfers toward major taxonomic hubs.

Countries with stronger environmental governance exhibited lower holotype outflow but greater appropriation of foreign holotypes, suggesting that governance shapes both the retention and accumulation of taxonomic resources. This pattern may reflect greater institutional capacity to regulate biological resources and sustain scientific collections over time, consistent with arguments that environmental legislation is most effective when backed by institutions capable of implementing its biological goals (Karr, 1990). At finer scales, the probability of local holotype retention increased for specimens collected within protected areas (61.7% versus 53.2%; Fig. 5d), suggesting that permitting systems and institutional oversight may enhance the traceability and accountability of biological collections (Cisneros et al., 2022; SNUC, 2000). Nevertheless, this contrasts with concerns that biodiversity regulations inspired by the Convention on Biological Diversity can inadvertently hinder taxonomic research and specimen exchange when implemented through overly restrictive access frameworks (Prathapan et al., 2018; Prathapan & Rajan, 2020), suggesting that the effects of governance depend on how conservation objectives are balanced with scientific access and capacity building.

Holotype appropriation was also greater in countries with higher research investment and stronger colonial legacies, mirroring drivers associated with other biodiversity knowledge shortfalls (Guedes et al., 2023; C. Meyer et al., 2015; Titley et al., 2017). These findings indicate that the same factors enabling biodiversity knowledge production also helps to concentrate the material resources upon which that knowledge depends. Although a greater number of research institutions promotes local holotype retention, it also facilitates holotype appropriation, illustrating how the same factor can simultaneously operate in competing push-pull processes (Fig. 5c). Repository countries are generally wealthier and possess substantially greater scientific infrastructure, suggesting that although retention and appropriation share similar drivers, these processes operate across different levels of scientific capacity (Fig. S10). Indeed, bilateral transfers of holotypes were better explained by differences between countries than by the characteristics of origin or destination countries alone (Fig. 5b), highlighting the importance of disparities in scientific capacity. High investments in biodiversity science increase a country’s capacity to study its own but also foreign biodiversity. This is exacerbated by salary inequalities that even allow self-funded researchers to study biodiversity hotspots even when local experts cannot afford it (Farooq & Nganhane, 2026).

Our findings reveal that the global distribution of taxonomic reference material emerges from the interaction of three distinct but interconnected processes: retention, extraction, and appropriation. While local scientific capacity and governance favour the retention of holotypes near their regions of origin, biodiversity-rich areas remain vulnerable to extraction, and foreign holotype accumulation is concentrated in a limited set of scientifically and historically advantaged nations. These patterns indicate that reducing inequalities in biodiversity documentation requires more than increasing species discovery alone; it also requires addressing where the material foundations of taxonomic knowledge are ultimately housed. Recent proposals such as specimen deposition plans, regional retention policies, co-curation agreements, and improved access to collections offer practical opportunities to increase transparency and equity in holotype stewardship (Moura et al., 2026). More fundamentally, reducing geographic inequalities in holotype stewardship will likely require sustained investments in taxonomic capacity building, as exemplified by long-term initiatives that strengthen expertise, collections, and institutional infrastructure (Rodman & Cody, 2003). By identifying the drivers underlying retention, extraction, and appropriation, our study provides a framework for understanding how taxonomic resources become geographically concentrated and for guiding efforts toward a more equitable global distribution of biodiversity knowledge and its material foundations.

## Acknowledgements

Our sincere gratitude is extended to the global community of taxonomists, field naturalists, and scientists, past and present, whose dedicated work in discovering and describing new reptile species has made this study possible.

## Author contributions

MRM conceived the study, all authors compiled the data, MRM analysed the data, MRM developed the figures, MRM and MTM wrote the text. All authors provided critical feedback and helped shape the research, analysis, and manuscript.

## Data availability

The data to reproduce the results of this study will be made available via Zenodo Digital Repository.

## Code availability

The code for reproducing the results of this study will be made available at Zenodo Digital Repository.

## Additional information

Supplementary Information is available for this manuscript, including Supplementary Tables S1 to S6 and Supplementary Figures S1 to S10.

