## Supplementary Material for "Biodiversity material remains where capacity and governance are strong, but taxonomic resources concentrate with geopolitical power"

**This PDF file includes**

Supplementary Tables S1-S6

Supplementary Figures S1-S10

### SUPPLEMENTARY TABLES

**Table S1. Variation Inflation Factor (VIF) for predictor variables of holotype retention and export counts (1990-2024).** Analyses performed at the species-level included all predictors, whereas analyses performed at the level of country-pairs (bilateral export counts) included only country-level predictors.

| **Variable** | **Species-level dataset** | **Bilateral flows based on origin countries** | **Bilateral flows based on destination countries** | **Bilateral flows based on disparity values** |
|---|---|---|---|---|
| Socioeconomic interest | 1.11 | | | |
| Year of description | 1.46 | | | |
| Number of authors | 1.49 | | | |
| Collector inclusion | 1.08 | | | |
| Collection site protection status | 1.02 | | | |
| Taxonomic revision | 1.03 | | | |
| Maximum body mass | 1.12 | | | |
| Absolute latitude | 1.24 | | | |
| *Colonial origin | 1.81 | 1.25 | 1.06 | 1.45 |
| *Research spending (% of GDP) | 3.16 | 2.87 | 3.85 | 2.02 |
| *Researchers per capita | 4.39 | 3.47 | ** | 2.9 |
| *Number of collected specimens | 2.34 | 1.97 | 4.29 | 2.1 |
| *Number of research institutions | 3.35 | 7.17 | 2.98 | 6.23 |
| *Average endemism richness | 2.08 | 3.62 | 7.08 | 3.38 |
| *Biodiversity and Habitat Issue | 1.15 | 1.3 | 1.55 | 1.19 |
| *Global Peace Index | 1.27 | 1.32 | 2.07 | 1.44 |
| *Human Capital Index | 2.35 | 3.32 | 2.85 | 2.43 |

* Country-level predictors (national indicators).

** Variable removed following the multicollinearity assessment procedure.

**Table S2. Variation Inflation Factor (VIF) for predictor variables of country-level analyses (1990-2024).** Analyses of holotype retention and outflow centrality used information on origin-countries, while analyses of holotype appropriation and inflow centrality used data on destination-countries.

| **Variable** | **Per-country holotype retention** | **Per-country outflow centrality** |
|---|---|---|
| ***Models using origin-countries*** | | |
| Colonial origin | 1.86 | 1.68 |
| Research spending (% of GDP) | 3.48 | 2.24 |
| Researchers per capita | 4.02 | 3.24 |
| Number of collected specimens | 2.08 | 2.02 |
| Number of research institutions | 3.99 | 5.02 |
| Average endemism richness | 2.71 | 2.66 |
| Biodiversity and Habitat Issue | 1.19 | 1.18 |
| Global Peace Index | 1.34 | 1.3 |
| Human Capital Index | 2.52 | 2.67 |
| **Variable** | **Per-country holotype appropriation** | **Per-country inflow centrality** |
| ***Models using destination-countries*** | | |
| Colonial origin | 3.84 | 2.4 |
| Research spending (% of GDP) | 5.02 | 5.58 |
| Researchers per capita | ** | ** |
| Number of collected specimens | 4.34 | 3.07 |
| Number of research institutions | 4.51 | 4.65 |
| Average endemism richness | 6.7 | 4.25 |
| Biodiversity and Habitat Issue | 1.51 | 1.41 |
| Global Peace Index | 1.62 | 1.52 |
| Human Capital Index | 3.82 | 3.41 |

** Variable removed following the multicollinearity assessment procedure.

**Table S3. Standardized coefficients from the species-level generalized linear mixed model (binomial GLMM) evaluating predictors of holotype retention probability (1990-2024).** Odds ratios (OR), 95% confidence intervals (CI), and p-values from the species-level generalized linear mixed model (binomial GLMM) evaluating predictors of holotype retention probability. Odds ratios greater than 1 indicate increased likelihood of local holotype retention, whereas values below 1 indicate reduced retention probability. The model included country of origin as a random effect. Significant predictors ($p < 0.05$) are highlighted in bold.

| Variable | Odds Ratio | Lower CI | Upper CI | p-value |
|---|---|---|---|---|
| Socioeconomic interest | 1.360 | 1.045 | 1.770 | **0.022** |
| Absolute latitude | 1.024 | 0.795 | 1.320 | 0.852 |
| Year of description | 1.846 | 1.601 | 2.128 | **<0.001** |
| Collector inclusion | 2.804 | 2.132 | 3.686 | **<0.001** |
| Taxonomic revision | 0.496 | 0.382 | 0.644 | **<0.001** |
| Collection site protection status | 1.416 | 1.048 | 1.915 | **0.024** |
| Maximum body mass | 0.867 | 0.772 | 0.974 | **0.016** |
| Number of authors | 1.269 | 1.102 | 1.460 | **0.001** |
| *Average endemism richness | 1.414 | 0.507 | 3.946 | 0.508 |
| *Biodiversity and Habitat Issue | 1.033 | 0.585 | 1.827 | 0.910 |
| *Colonial origin | 1.889 | 0.433 | 8.236 | 0.397 |
| *Global Peace Index | 1.049 | 0.625 | 1.760 | 0.855 |
| *Human Capital Index | 4.660 | 2.037 | 10.661 | **<0.001** |
| *Number of collected specimens | 0.536 | 0.259 | 1.110 | 0.093 |
| *Number of research institutions | 14.130 | 5.085 | 39.267 | **<0.001** |
| *Research spending (% of GDP) | 0.832 | 0.351 | 1.974 | 0.677 |
| *Researchers per capita | 0.948 | 0.360 | 2.497 | 0.914 |

* Country-level predictors (national indicators).

**Table S4. Standardized coefficients from the generalized linear mixed model evaluating predictors of bilateral holotype export counts between country pairs (1990-2024).** Holotype export counts were modelled using origin-, destination-, or intercountry-differences (Δ = destination value − origin value) predictors. Results are presented as model coefficients (β ± SE), 95% confidence intervals (CI), and p-values The model included country of origin as a random effect. Significant predictors ($p < 0.05$) are highlighted in bold.

| **Country-level attribute** | **Estimate (β)** | **SE** | **Lower CI** | **Upper CI** | **p-value** |
|---|---|---|---|---|---|
| ***Origin-country*** | | | | | |
| Average endemism richness | 1.495 | 0.250 | 1.006 | 1.985 | **<0.001** |
| Biodiversity and Habitat Issue | -0.254 | 0.132 | -0.512 | 0.004 | 0.053 |
| Colonial origin | 0.390 | 0.380 | -0.356 | 1.135 | 0.305 |
| Global Peace Index | -0.342 | 0.124 | -0.585 | -0.099 | **0.006** |
| Human Capital Index | -0.026 | 0.219 | -0.454 | 0.402 | 0.906 |
| Number of collected specimens | 0.408 | 0.187 | 0.043 | 0.774 | **0.029** |
| Number of research institutions | -0.728 | 0.325 | -1.365 | -0.091 | **0.025** |
| Research spending (% of GDP) | -0.311 | 0.192 | -0.686 | 0.065 | 0.105 |
| Researchers per capita | 0.150 | 0.203 | -0.248 | 0.548 | 0.461 |
| ***Destination-country*** | | | | | |
| Average endemism richness | 0.009 | 0.610 | -1.187 | 1.205 | 0.988 |
| Biodiversity and Habitat Issue | 0.103 | 0.304 | -0.493 | 0.699 | 0.735 |
| Colonial origin | 0.340 | 0.417 | -0.478 | 1.158 | 0.415 |
| Global Peace Index | 0.134 | 0.356 | -0.563 | 0.831 | 0.707 |
| Human Capital Index | 0.022 | 0.319 | -0.604 | 0.648 | 0.945 |
| Number of collected specimens | 0.736 | 0.623 | -0.485 | 1.957 | 0.237 |
| Number of research institutions | -0.123 | 0.434 | -0.973 | 0.728 | 0.778 |
| Research spending (% of GDP) | 0.251 | 0.411 | -0.554 | 1.056 | 0.541 |
| ***Intercountry differences (Δ = destination value – origin value)*** | | | | | |
| Average endemism richness | -2.878 | 0.532 | -3.922 | -1.835 | **<0.001** |
| Biodiversity and Habitat Issue | 0.057 | 0.188 | -0.311 | 0.426 | 0.761 |
| Colonial origin | 0.300 | 0.380 | -0.445 | 1.044 | 0.430 |
| Global Peace Index | 0.802 | 0.219 | 0.372 | 1.231 | **<0.001** |
| Human Capital Index | 0.264 | 0.218 | -0.164 | 0.692 | 0.226 |
| Number of collected specimens | -0.054 | 0.375 | -0.790 | 0.682 | 0.886 |
| Number of research institutions | 0.899 | 0.452 | 0.013 | 1.784 | **0.047** |
| Research spending (% of GDP) | 0.116 | 0.211 | -0.296 | 0.529 | 0.581 |
| Researchers per capita | -0.204 | 0.235 | -0.665 | 0.256 | 0.385 |

**Table S5. Standardized coefficients from country-level generalized linear models evaluating predictors of holotype retention and holotype appropriation (1990-2024).** Holotype retention was modelled as counts of retained versus exported holotypes conditional on the total number of holotypes associated with each country using a beta-binomial distribution, whereas holotype appropriation was modelled as counts of imported holotypes conditional on collection size (total housed species) using a negative binomial distribution with a log-transformed offset term. Results are presented as standardized coefficients (β ± SE), 95% confidence intervals (CI), and p-values. Significant predictors ($p < 0.05$) are shown in bold.

| Country-level attribute | Estimate (β) | SE | Lower CI | Upper CI | p-value |
|---|---|---|---|---|---|
| ***Holotype retention*** | | | | | |
| Average endemism richness | 0.279 | 0.312 | -0.332 | 0.890 | 0.370 |
| Biodiversity and Habitat Issue | -0.242 | 0.176 | -0.588 | 0.103 | 0.170 |
| Colonial origin | 0.090 | 0.495 | -0.881 | 1.060 | 0.856 |
| Global Peace Index | 0.012 | 0.182 | -0.345 | 0.369 | 0.948 |
| Human Capital Index | 1.507 | 0.368 | 0.787 | 2.228 | **<0.001** |
| Number of collected specimens | -0.424 | 0.223 | -0.862 | 0.013 | 0.057 |
| Number of research institutions | 1.293 | 0.377 | 0.553 | 2.032 | **0.001** |
| Research spending (% of GDP) | -0.063 | 0.299 | -0.650 | 0.524 | 0.833 |
| Researchers per capita | -0.153 | 0.318 | -0.777 | 0.470 | 0.629 |
| ***Holotype appropriation*** | | | | | |
| Average endemism richness | -0.380 | 0.435 | -1.232 | 0.473 | 0.383 |
| Biodiversity and Habitat Issue | 0.470 | 0.206 | 0.066 | 0.875 | **0.023** |
| Colonial origin | 0.083 | 0.653 | -1.197 | 1.363 | 0.898 |
| Global Peace Index | 0.308 | 0.193 | -0.072 | 0.687 | 0.112 |
| Human Capital Index | -0.206 | 0.426 | -1.040 | 0.629 | 0.629 |
| Number of collected specimens | -0.028 | 0.335 | -0.684 | 0.628 | 0.934 |
| Number of research institutions | -0.138 | 0.342 | -0.809 | 0.533 | 0.687 |
| Research spending (% of GDP) | 1.609 | 0.523 | 0.584 | 2.635 | **0.002** |

**Table S6. Standardized coefficients from country-level generalized linear models evaluating predictors of inflow and outflow centrality in the global holotype exchange network (1990-2024).** Inflow centrality (Katz in-centrality) represents the relative extractive influence of countries in acquiring foreign holotypes, whereas outflow centrality (Katz out-centrality) reflects the extent to which holotypes are exported to foreign countries. Both response variables were modelled using gamma distributions with log link functions. Results are presented as standardized coefficients (β ± SE), 95% confidence intervals (CI), and p-values. Significant predictors ($p < 0.05$) are shown in bold.

| **Country-level attribute** | **Estimate (β)** | **SE** | **Lower CI** | **Upper CI** | **p-value** |
|---|---|---|---|---|---|
| ***Outflow centrality*** | | | | | |
| Average endemism richness | 0.024 | 0.004 | 0.017 | 0.031 | **<0.001** |
| Biodiversity and Habitat Issue | -0.009 | 0.002 | -0.014 | -0.004 | **<0.001** |
| Colonial origin | 0.006 | 0.007 | -0.008 | 0.020 | 0.409 |
| Global Peace Index | -0.002 | 0.002 | -0.007 | 0.002 | 0.361 |
| Human Capital Index | -0.001 | 0.003 | -0.008 | 0.005 | 0.690 |
| Number of collected specimens | 0.000 | 0.003 | -0.006 | 0.006 | 0.935 |
| Number of research institutions | -0.005 | 0.005 | -0.015 | 0.004 | 0.251 |
| Research spending (% of GDP) | -0.002 | 0.003 | -0.008 | 0.004 | 0.514 |
| Researchers per capita | 0.001 | 0.004 | -0.006 | 0.009 | 0.690 |
| ***Inflow centrality*** | | | | | |
| Average endemism richness | -0.348 | 0.180 | -0.701 | 0.006 | 0.054 |
| Biodiversity and Habitat Issue | 0.254 | 0.099 | 0.060 | 0.448 | **0.010** |
| Colonial origin | 0.084 | 0.264 | -0.433 | 0.602 | 0.749 |
| Global Peace Index | 0.230 | 0.110 | 0.014 | 0.447 | **0.037** |
| Human Capital Index | -0.157 | 0.153 | -0.456 | 0.142 | 0.303 |
| Number of collected specimens | -0.049 | 0.129 | -0.301 | 0.203 | 0.705 |
| Number of research institutions | 0.679 | 0.157 | 0.371 | 0.987 | **<0.001** |
| Research spending (% of GDP) | 0.326 | 0.191 | -0.049 | 0.700 | 0.088 |
| Average endemism richness | -0.348 | 0.180 | -0.701 | 0.006 | 0.054 |

### SUPPLEMENTARY FIGURES

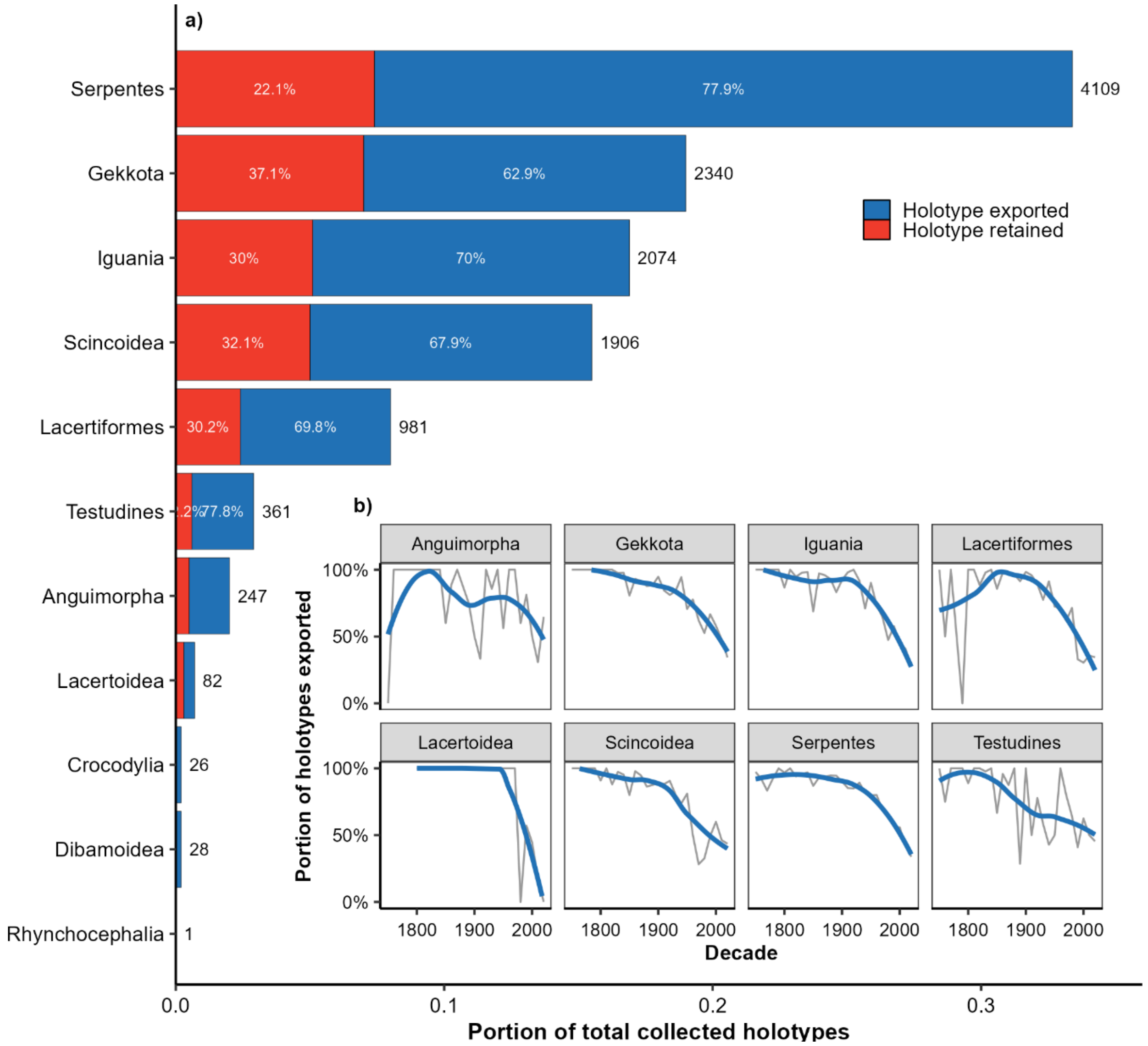

**Figure S1. Relative count of reptile holotype specimens across major clades.** (a) Proportion of reptile holotypes retained locally versus exported for the period between 1758 and 2024 across major clades. Numbers adjacent to bars indicate absolute values. (b) Proportion of species described in each decade whose holotypes were deposited outside their country of origin, with smoothed trajectories illustrating long-term shifts in specimen circulation. Only clades with more than 30 species described are show in the inset temporal plots.

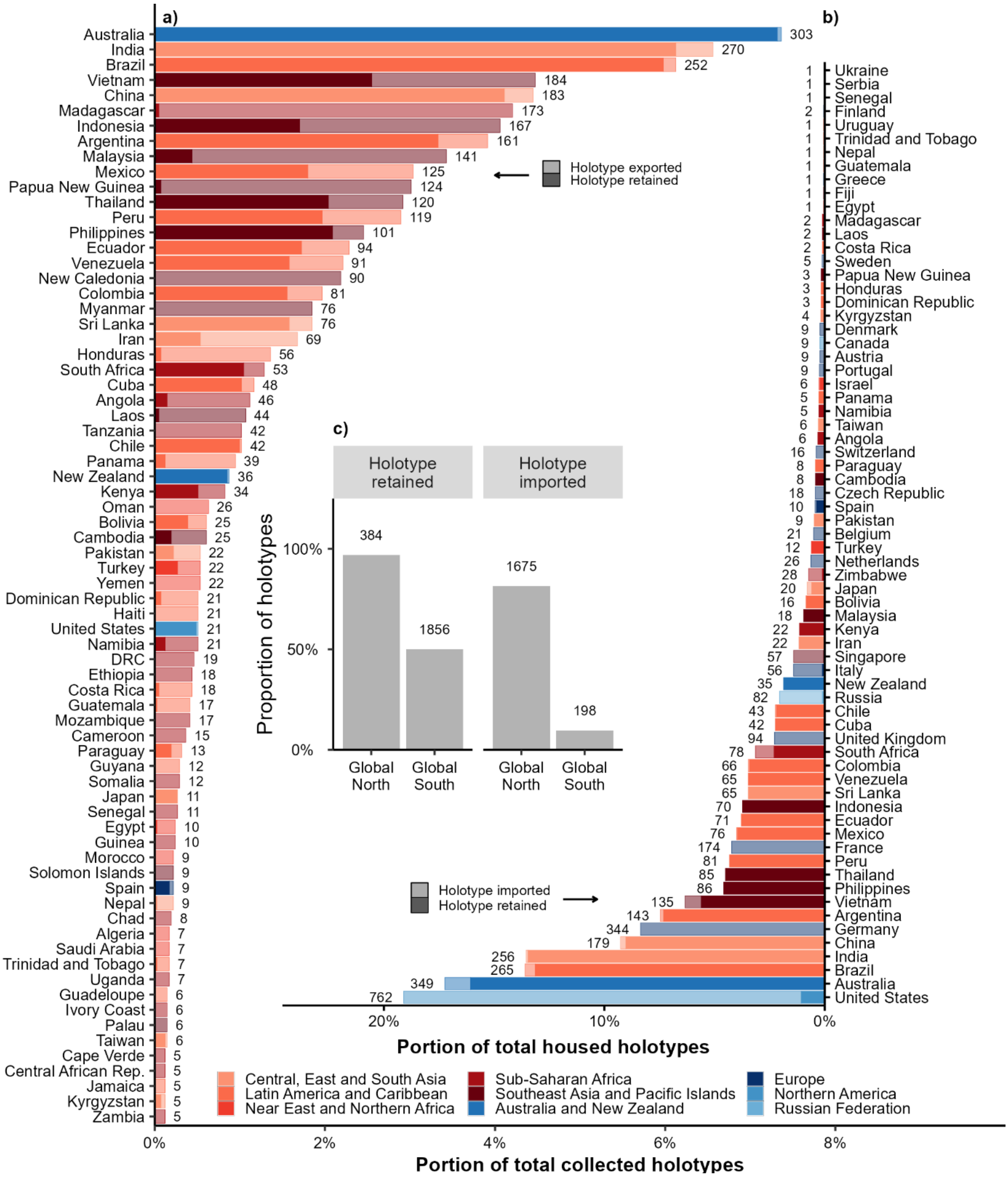

**Figure S2. Relative count of reptile holotype specimens across origin and destination countries (1990-2024).** Barplots show the per-country proportions of holotypes (a) retained locally versus exported and (b) proportions of holotypes imported versus retained. (c) Relative retention, exportation, and importation between the Global North and Global South. Numbers adjacent to bars indicate absolute values. Countries with <5 described species were omitted from the left main panel.

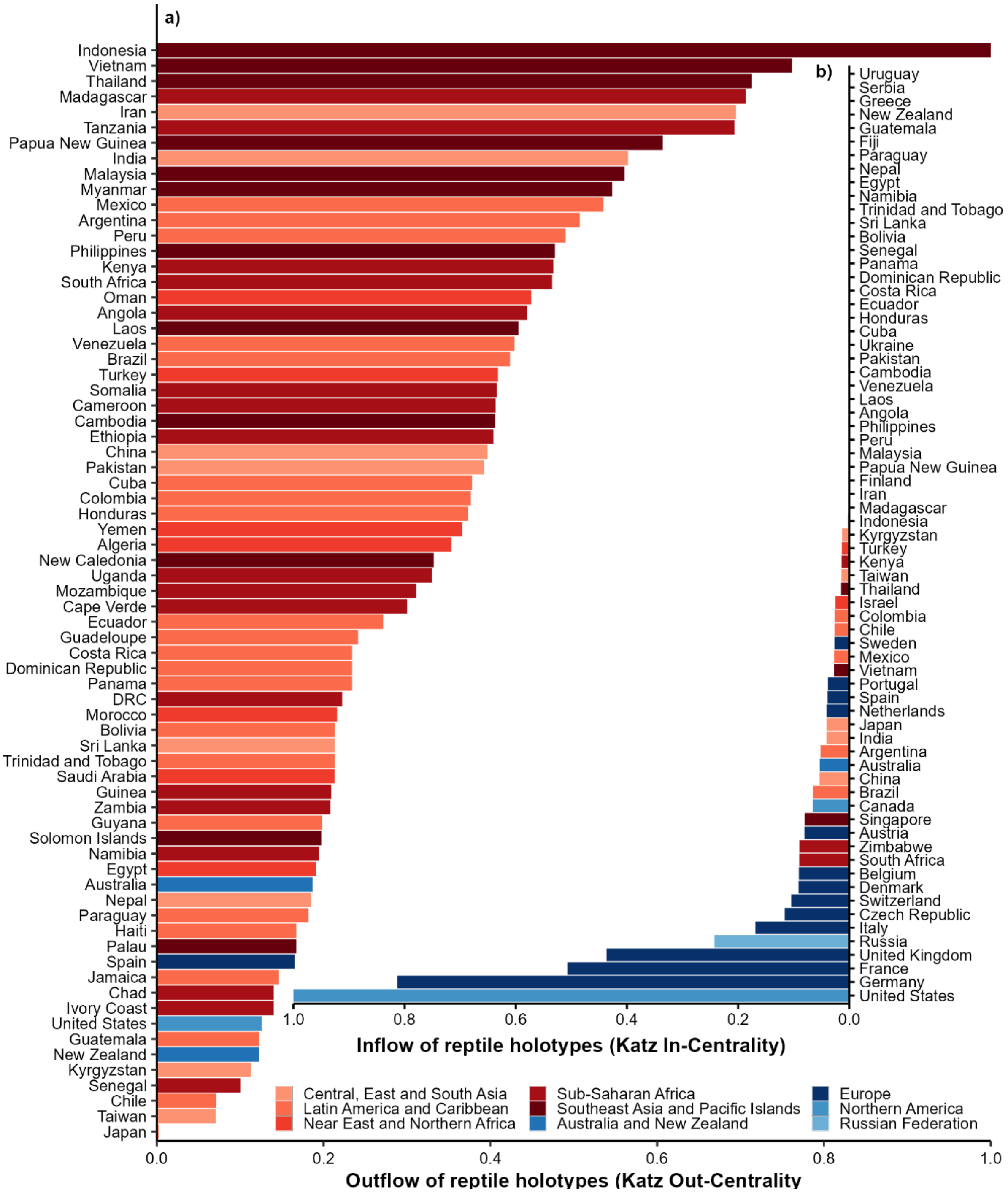

**Figure S3. Country-level inflow and outflow centrality of reptile holotype specimens (1990-2024).** Katz centrality metrics were rescaled to range from 0 to 1 to improve visualizaion. (a) Outflow centrality, only countries with ≥5 described species are shown. (b) Inflow centrality. High inflow centrality indicates countries exerting greater extractive influence, whereas high outflow centrality reflects countries more strongly subject to extraction.

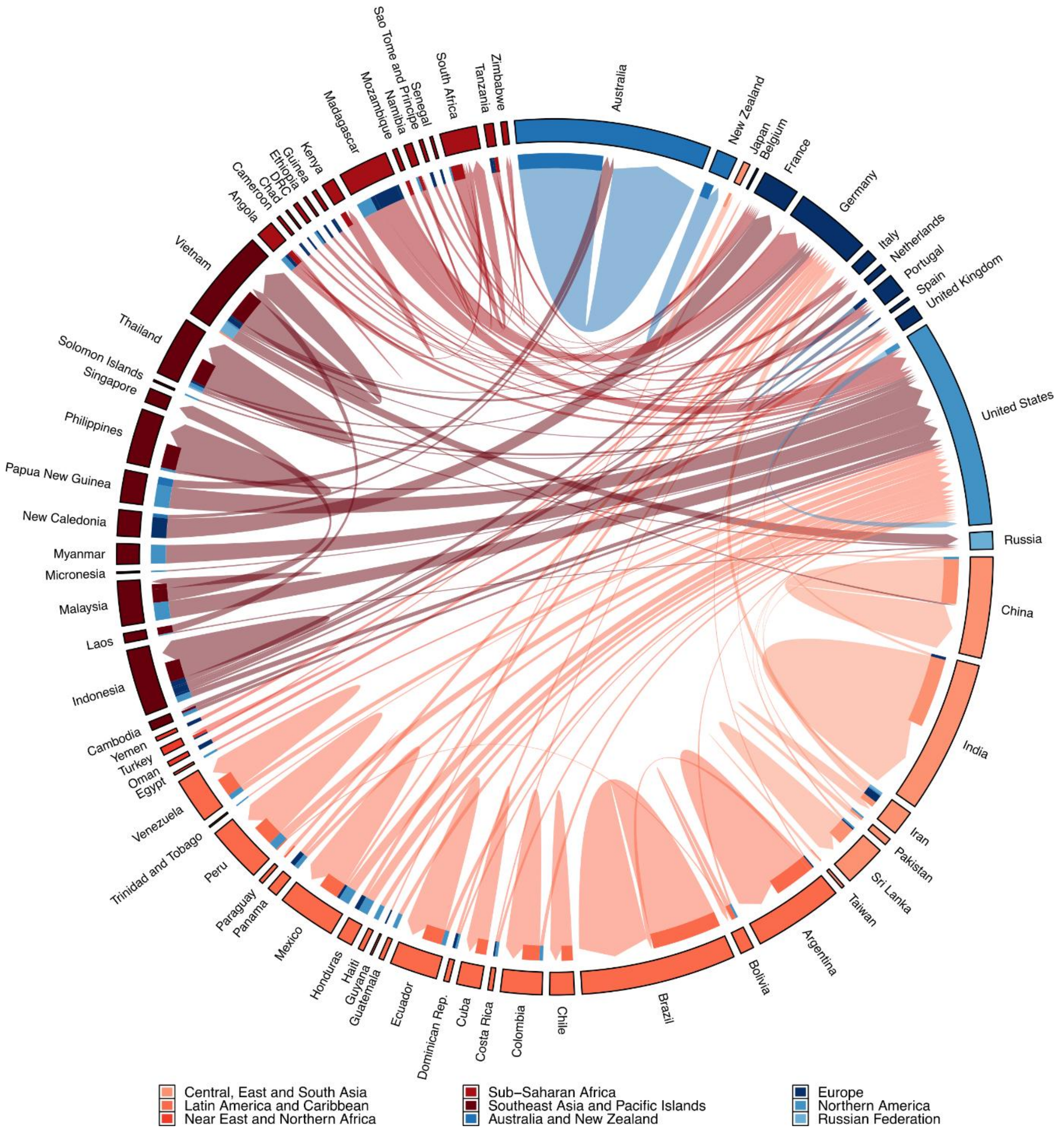

**Figure S4. Global flow of reptile holotype specimens across countries (1990-2024).** Only countries with at least five holotypes housed in foreign collections are shown. Arrows indicate source-to-repository pathways, with origin countries pointing to host nations.

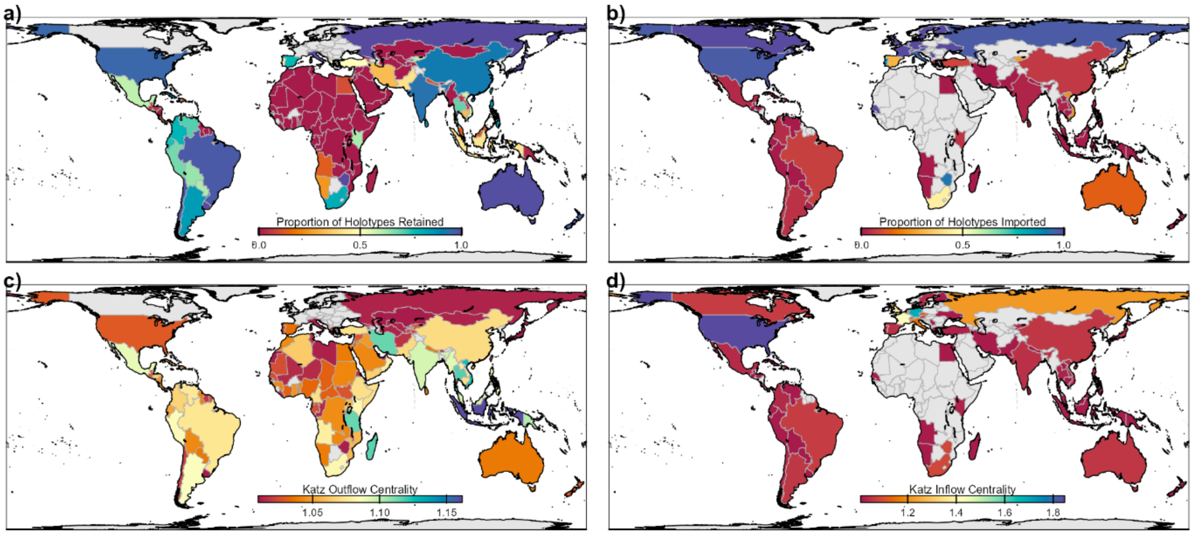

**Figure S5. Geographical patterns of reptile holotype transfer across countries (1990-2024)**. (a) Proportion of holotypes retained within the country of origin. (b) Proportion of housed holotypes that were imported from other countries. (c) Outflow centrality metric, representing the extent to which a country's holotypes are described by foreign institutions. (d) Inflow centrality metric, representing a country's role in describing holotypes originating elsewhere. Countries shown in light grey were neither origins (a, c) nor destinations (b, d) of reptile holotypes described in time investigated time period. All maps are presented using an equal-area projection.

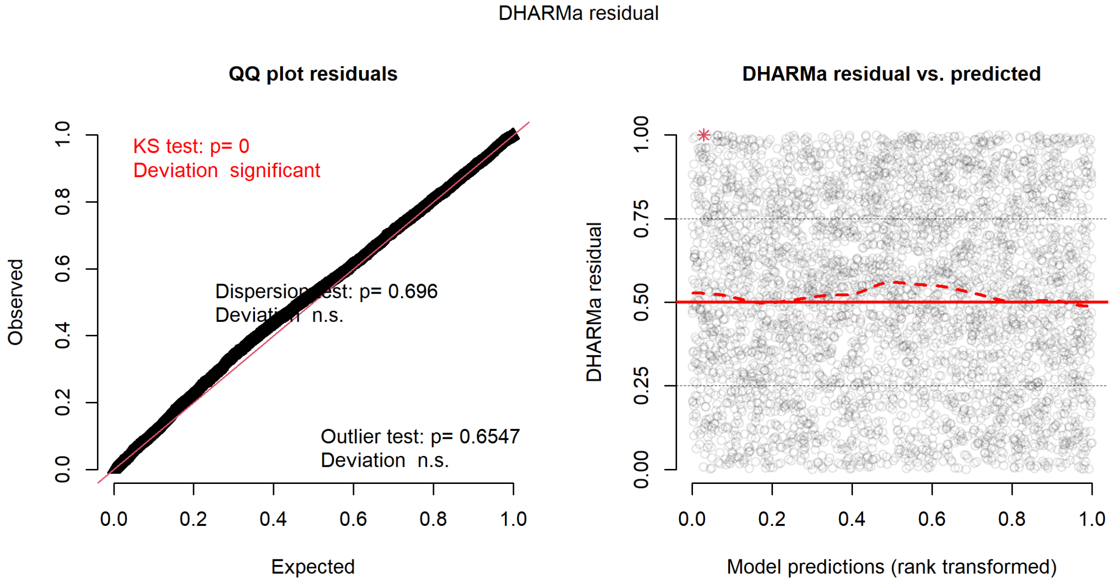

**Figure S6. DHARMa residual diagnostics for the species-level mixed model of holotype retention (1990-2024).** (a) Q-Q plot of scaled residuals against uniform distribution. (b) Residuals plotted against model predictions. A significant deviation in one diagnostic test does not necessarily invalidate our results when considered alongside others. The QQ plots indicated a generally good fit, and residual plots showed only minor deviations in one or two quantiles. This is not a major concern, particularly given our reasonably sample size.

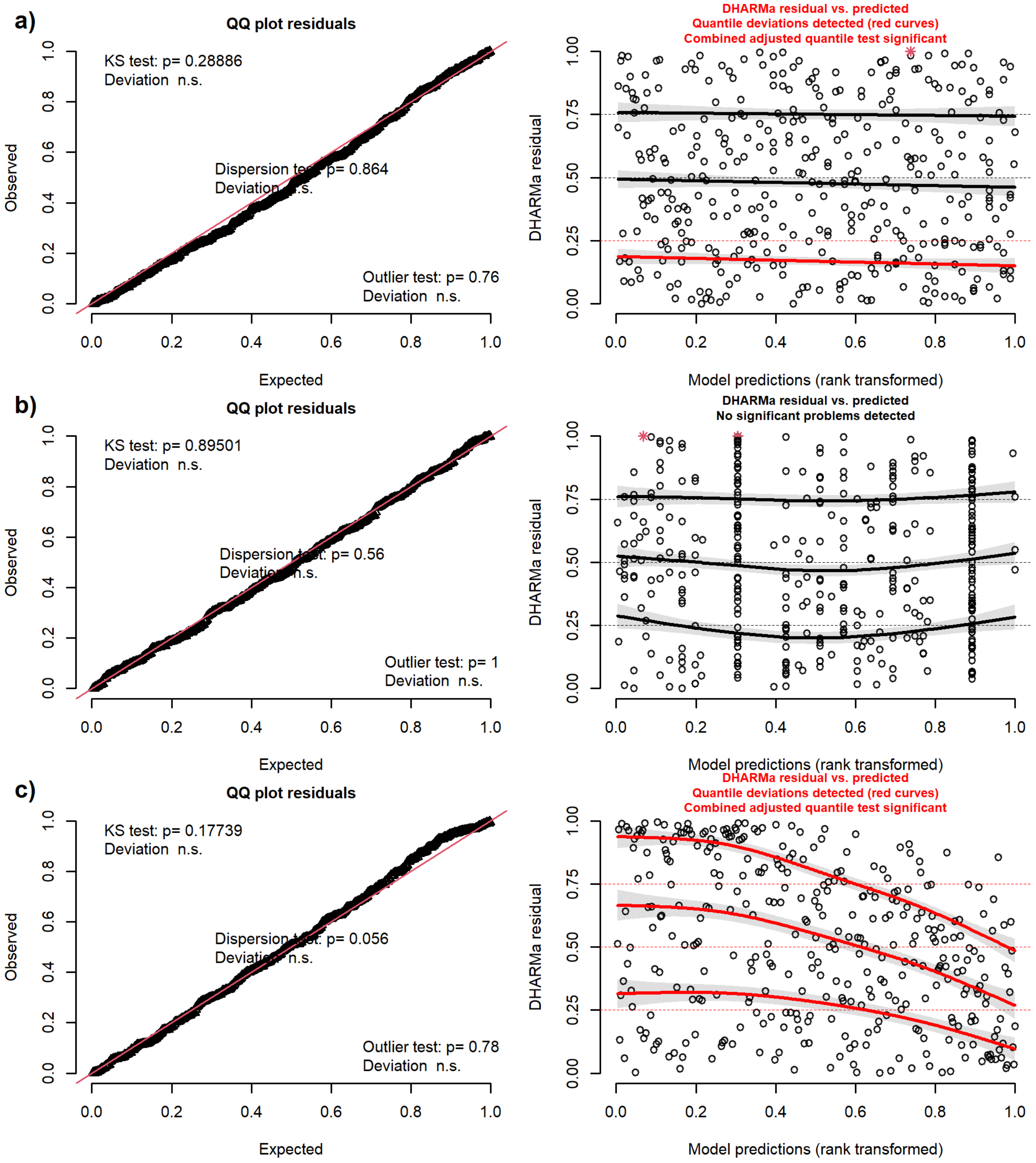

**Figure S7. DHARMa residual diagnostics for the mixed model of holotype export counts at the country-pair level (1990-2024).** Holotype export counts were modelled using (a) origin-country predictors, (b) destination-country predictors, and (c) country-disparity predictors (Δ = destination value − origin value). In each row, the left panel shows the Q-Q plot of scaled residuals against the expected uniform distribution, and the right panel shows residuals versus model predictions. Overall, diagnostics indicated a good model fit, with only minor deviations in one or two quantiles that are unlikely to affect inference given the large sample size.

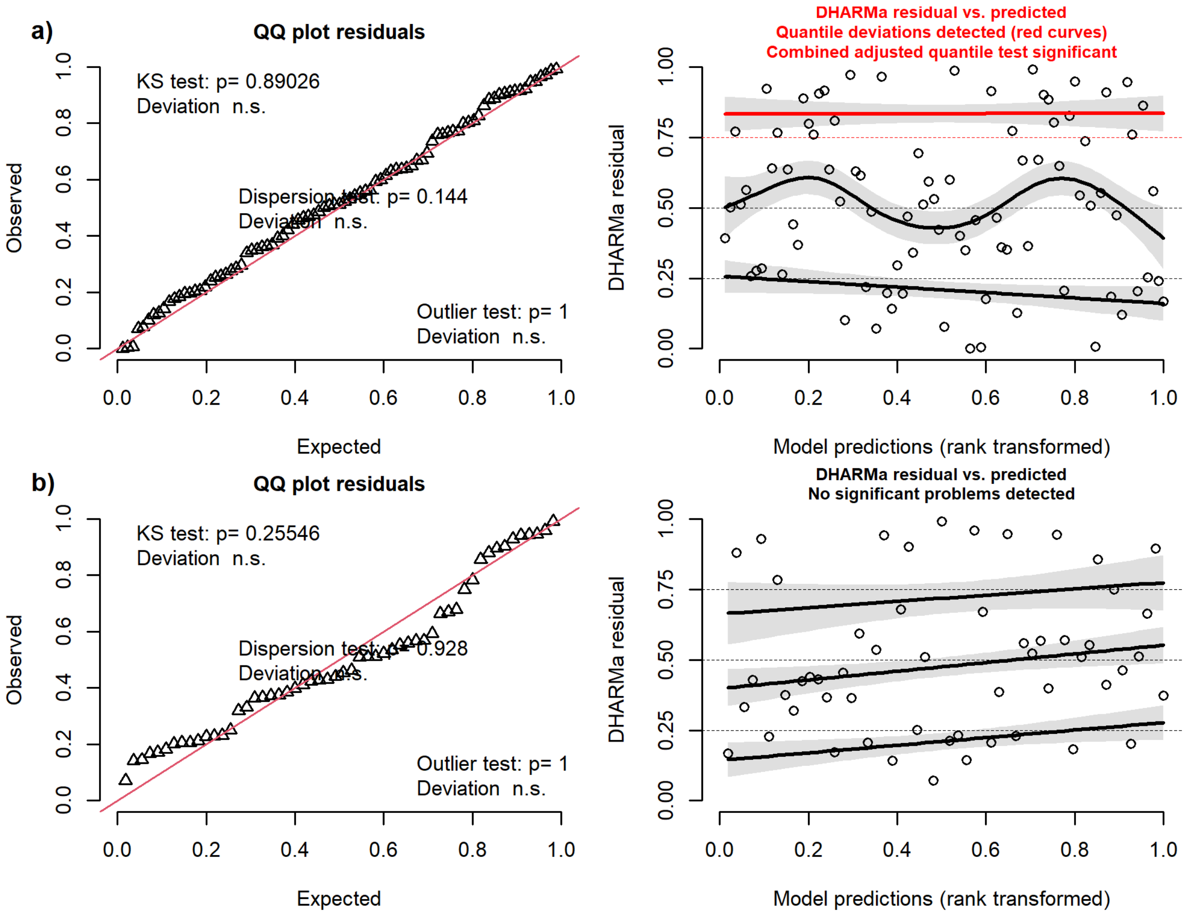

**Figure S8. DHARMa residual diagnostics for the model of holotype retention and appropriation at the country-level (1990-2024)**. (a) Holotype retention, modelled as counts of retained versus exported holotypes conditional on the total number of holotypes associated with each country. (b) Holotype appropriation, modelled as counts of imported holotypes conditional on collection size (total housed species). In each row, the left panel shows the Q-Q plot of scaled residuals against the expected uniform distribution, and the right panel shows residuals versus model predictions.

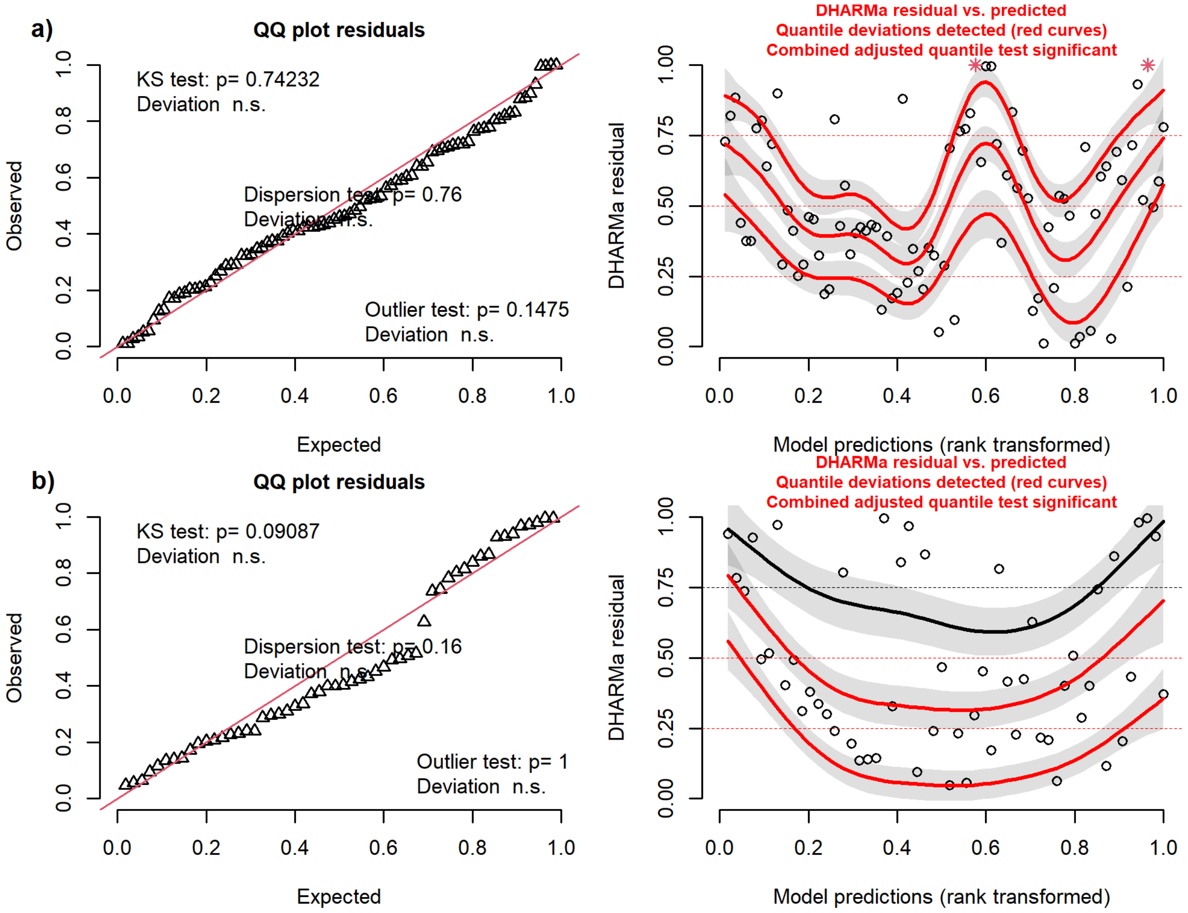

**Figure S9. DHARMa residual diagnostics for country-level models of inflow and outflow centrality in the global holotype exchange network (1990-2024).** (a) Inflow centrality (Katz in-centrality), quantified using Katz centrality based on incoming holotype exchanges. (b) Outflow centrality (Katz out-centrality), quantified using Katz centrality based on outgoing holotype exchanges. Both metrics were modelled using gamma distributions with log link functions. In each row, the left panel shows the Q-Q plot of scaled residuals against the expected uniform distribution, and the right panel shows residuals versus model predictions.

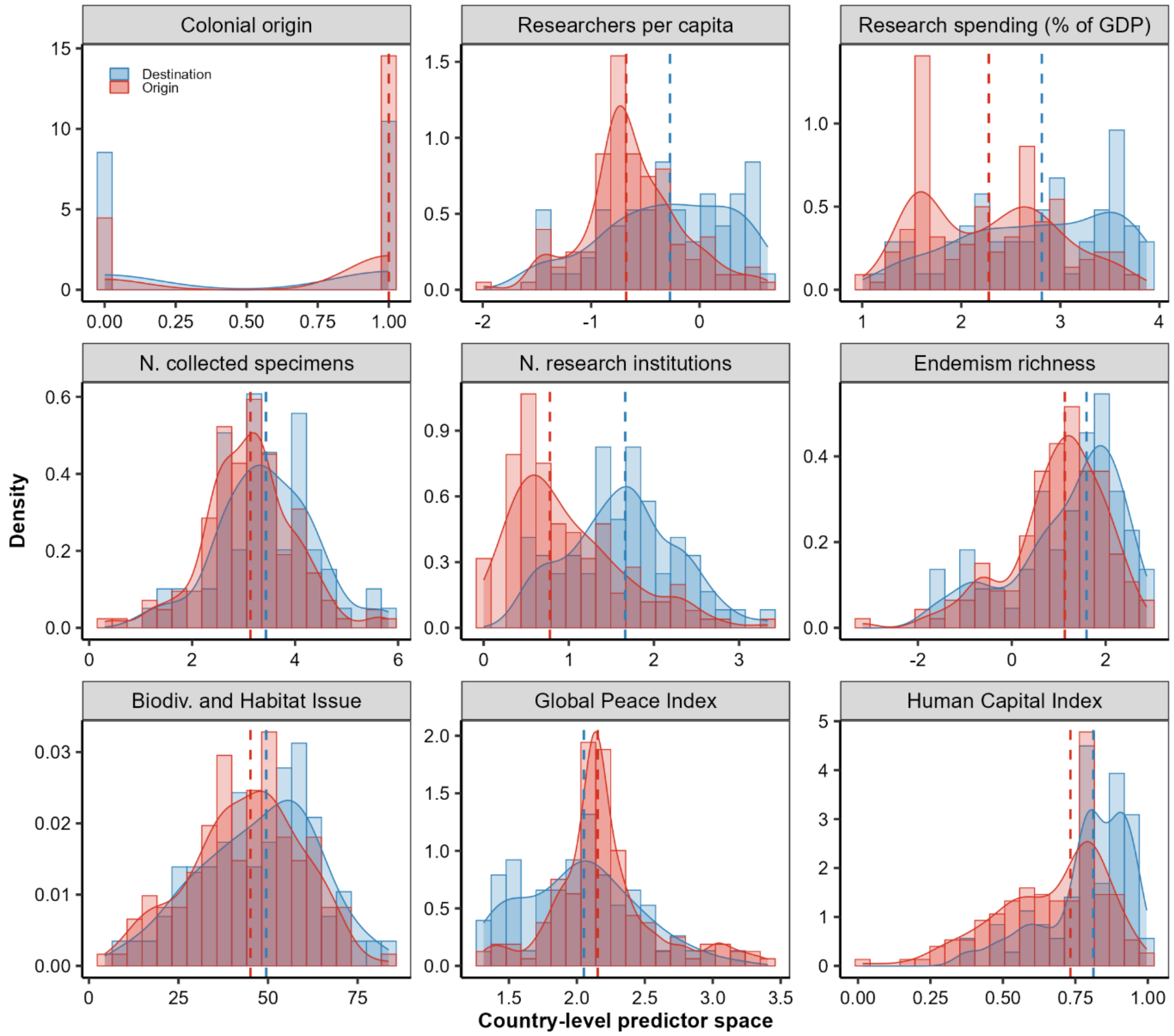

**Figure S10. Coverage of country-level predictor space by origin and destination countries in the global reptile holotype network.** Histograms and kernel density estimates depict the distributions of country-level predictors for origin- and destination-countries of reptile holotypes. Dashed vertical lines indicate median values for each group. Continuous variables describing researchers per capita, research spending (% of GDP), number of collected specimens, number of research institutions, and endemism richness were $\log_{10}$-transformed. The degree of overlap between distributions indicates the extent to which source and destination countries represent similar or distinct regions of the global predictor space.
